# FhaA plays a key role in mycobacterial polar elongation and asymmetric growth

**DOI:** 10.1101/2024.08.13.607006

**Authors:** Jessica Rossello, Bernardina Rivera, Maximiliano Anzibar Fialho, Ingrid Augusto, Magdalena Gil, Marina Andrea Forrellad, Fabiana Bigi, Azalia Rodríguez Taño, Estefanía Urdániz, Mariana Piuri, Kildare Miranda, Anne Marie Wehenkel, Pedro M Alzari, Leonel Malacrida, Rosario Durán

**Author notes:** Proteomic Platform, Mass Spectrometry for Biology Unit, CNRS UAR 2024, Institut Pasteur, Université Paris Cité, 75015, Paris, France.

## Abstract

Mycobacteria, including pathogens like *Mycobacterium tuberculosis*, exhibit unique growth patterns and cell envelope structures that challenge our understanding of bacterial physiology. This study sheds light on FhaA, a conserved protein in *Mycobacteriales*, revealing its pivotal role in coordinating cell envelope biogenesis and asymmetric growth.

The elucidation of the FhaA interactome in living mycobacterial cells reveals its participation in the protein network orchestrating cell envelope biogenesis and cell elongation/division. By manipulating FhaA levels, we uncovered its influence on cell morphology, cell envelope organization, and the localization of peptidoglycan biosynthesis machinery. Notably, *fhaA* deletion disrupted the characteristic asymmetric growth of mycobacteria, highlighting its importance in maintaining this distinctive feature.

Our findings position FhaA as a key regulator in a complex protein network, orchestrating the asymmetric distribution and activity of cell envelope biosynthetic machinery. This work not only advances our understanding of mycobacterial growth mechanisms but also identifies FhaA as a potential target for future studies on cell envelope biogenesis and bacterial growth regulation. These insights into the fundamental biology of mycobacteria may pave the way for novel approaches to combat mycobacterial infections addressing the ongoing challenge of diseases like tuberculosis in global health.

## Introduction

Mycobacterium *tuberculosis*, the causative agent of tuberculosis, is among the deadliest human pathogens. According to the World Health Organization, tuberculosis ranked as the first cause of death from a single bacterial infectious agent worldwide (1).

One of the peculiarities of this bacillus lies in its cell growth and division modes, which differ significantly from those of well-studied rod-shaped bacteria, such as *Escherichia coli or Bacillus subtilis* (2). Mycobacteria need to synthesize a complex cell wall during growth and division. This distinctive structure, composed of peptidoglycan (PG) covalently attached to arabinogalactans esterified with mycolic acids, is relevant for conferring intrinsic antibiotic resistance and plays a major role in host-pathogen interactions and virulence (3,4). Moreover, unlike model bacilli that incorporate new cell wall material laterally, mycobacteria exhibit an asymmetric polar elongation mode in which the old pole inherited from the mother cell outpaces the newly formed pole in the rate of cell wall synthesis (2,5). This asymmetric growth pattern contributes to a diversified population in terms of size and antibiotic susceptibility (6). Furthermore, many well-characterized key members of the protein complexes guiding elongation (elongasome) and division (divisome) in *E. coli* and *B. subtilis* are absent among mycobacteria (2,7). Hence the molecular mechanisms underlying cell growth and division in these bacteria remain largely unknown. Nevertheless, an increasing number of mycobacterial-specific cell division and elongation protein candidates have started to be identified, including two ForkHead-Associated (FHA) domain-containing proteins, FhaA and FhaB, which specifically recognize phospho-Thr residues (8–10).

FhaA and FhaB are part of a highly conserved operon in *Mycobacteriales,* that also encodes two Shape, Elongation, Division and Sporulation (SEDS) genes (*rodA* and *pbpA*), two Ser/Thr protein kinases (*pknA* and *pknB*) and the unique phosphoserine/threonine protein phosphatase of the genome (11,12), pointing to its critical role in cell morphology, growth and its phospho-regulation.

Here, we focused on *M. tuberculosis* FhaA, a still poorly characterized multidomain protein. FhaA presents a C-terminal ForkHead-Associated (FHA) domain, which specifically recognizes phosphorylated Thr residues, linked by a ∼300 amino acid-long unstructured linker to an N-terminal globular domain with no similarity to any known protein (13). Previous reports provide evidence that supports a role for FhaA in the regulation of cell wall biosynthesis through its interaction with two phosphorylated PknB substrates. FhaA was proposed to inhibit the translocation of PG precursors from the cytosol to the periplasm through its interaction with the phosphorylated pseudokinase domain of the Lipid II flippase Mvin (10). It was also shown to interact with phosphorylated CwlM and potentially regulate the biosynthesis of PG precursors (14). In addition, knocking out *fhaA* in *Mycobacterium smegmatis* resulted in a short-cell phenotype (15), while its depletion caused increased accumulation of nascent PG at the poles and septa (10). Some of this previous data are difficult to reconcile, making the roles of FhaA and its molecular mechanisms still unclear.

Here, we explored protein-protein interactions involving mycobacterial FhaA in living cells. Our results showed that FhaA is part of an extensive protein network linking cell envelope biogenesis to cell elongation/division in mycobacteria. Overexpressing FhaA in *M. smegmatis* cells leads to alterations in composition and/or organization of the cell envelope along with mislocalization of the PG biosynthesis machinery, whereas deletion of the *fhaA* gene results in elongation defects and the loss of asymmetrical insertion of new cell wall material at the poles. Collectively, our findings indicate that FhaA plays a crucial role in polar growth by regulating the precise subcellular localization and asymmetric distribution of the cell envelope biosynthetic machinery organized within the elongasome.

## Results

### FhaA interactome in living cells

To decipher the FhaA interactome within live mycobacteria, we employed an unbiased methodology encompassing the overexpression of *M. tuberculosis* Strep-tagged FhaA in *M. smegmatis*. The method relied on a combination of chemical crosslinking, affinity purification and protein identification through MS (Figure 1A). Formaldehyde was selected as the crosslinking agent due to its ability to penetrate the highly hydrophobic cell envelope of mycobacteria and covalently link amino acids in close proximity (16). *M. smegmatis* transformed with the empty plasmid was used as control. To define the FhaA interactome, we compared the proteins recovered by affinity purification in control and *Msmeg_fhaA* strains, using in 5 biological replicates per condition. As shown in Figure 1B, 25 proteins were exclusively detected in at least 4 replicates of the purified protein complexes from *Msmeg_fhaA* (p≤0.01) (Table 1). The list includes: FhaA itself; Mvin, the flippase previously reported to interact with FhaA (10), and 23 putative direct or indirect FhaA interactors (Table 1 and Table S1). In addition, from 736 proteins identified in affinity purified fractions from both *Msmeg_fhaA* and control strains, 31 were statistically overrepresented in complexes isolated from *Msmeg_fhaA* (fold change ≥ 2; F-stringency: 0.04; q-value ≤ 0.05) (Figure 1C, Table S2). Altogether, we report a list of 55 proteins that represent putative direct or indirect FhaA interactors (Table 1 and Tables S1 and S2). Remarkably, the FhaA interactome comprises proteins with known and predicted physical and functional associations, unveiling statistically enriched compartments, which include the cell pole, cell tip, cell septum, and membrane fraction. (Table S3). Among the polar interactors (10,17–20), two of them (MSMEG_0317 and MSMEG_3080) exhibit an asymmetric distribution, specifically targeting the fast-growing pole (Table 1). Furthermore, functional enrichment analysis emphasized various interconnected biological processes, encompassing the regulation of developmental processes, cell shape regulation, as well as cell cycle and division regulation (Table S3). The recovery of previously known interactors, along with proteins that share the same subcellular localization and are involved in the same biological processes as FhaA, points to a reliable and physiologically relevant interactome.

**Figure 1.**
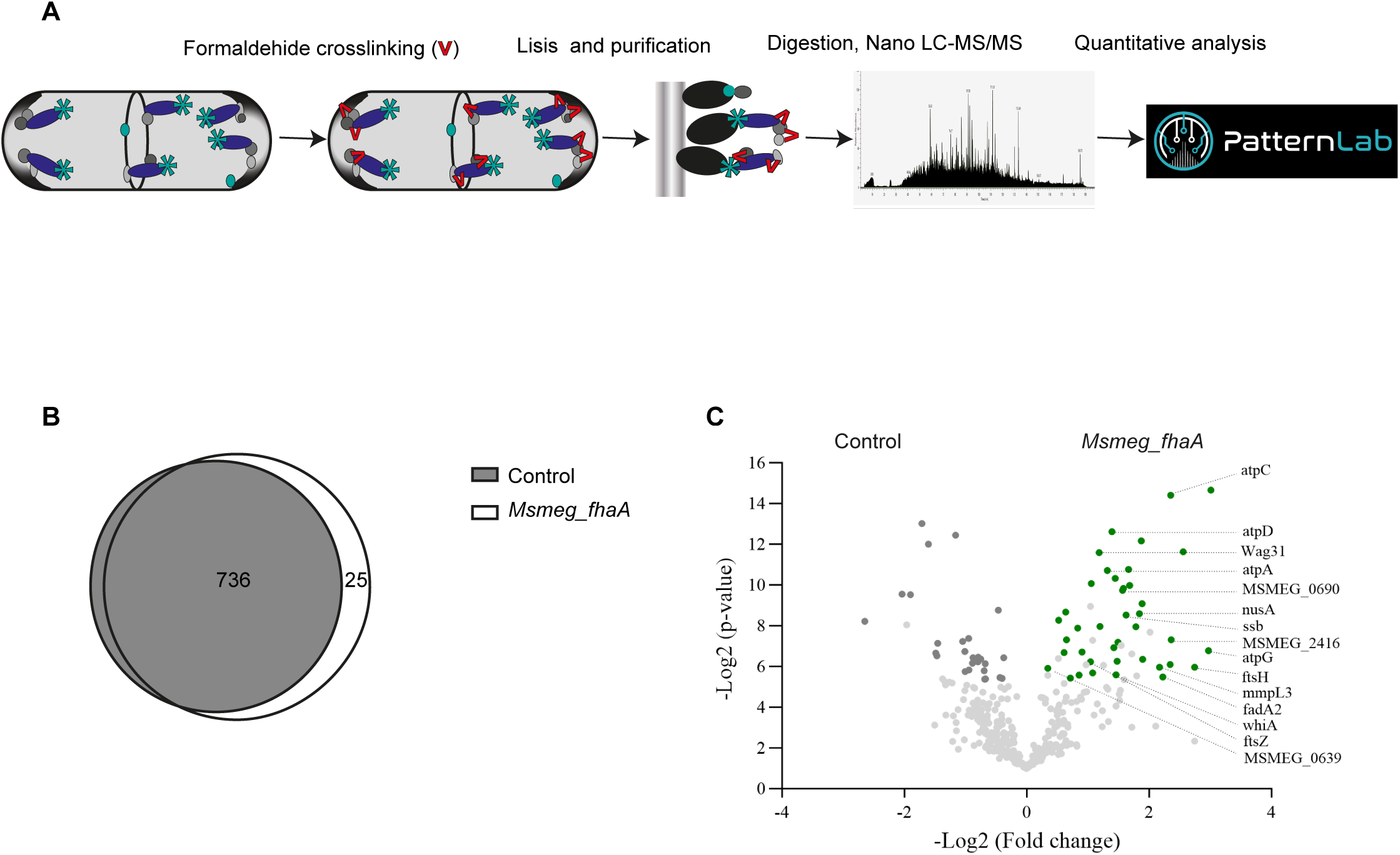
FhaA interactome in the living cell. **A)** Scheme of the strategy used to identify FhaA interacting proteins. Cultures of *M. smegmatis* overexpressing *M. tuberculosis* FhaA fused to Streptag were incubated with formaldehyde. FhaA covalently linked to its protein partners were purified using Strep-Tactin® columns and the recovered proteins were digested and identified by nano LC-MS/MS. **B)** Venn diagram showing the number of proteins identified in *Msmeg_fhaA* and control strains after affinity chromatography. Using the probability mode of Patternlab Venn diagram module, 25 proteins were statistically identified as exclusive of FhaA interactome (p <0.01). (Table 1 and Table S1). **C)** Volcano plot showing proteins identified in at least 7 replicates of the 10 replicates analysed, plotted according to its p-value (log2 p) and fold change (log2 Fold change). Proteins statistically enriched in FhaA complexes (q-value≤0.05) with a fold change greater than 2 are displayed in green, and those related to cell elongation/cell envelope biosynthesis are labelled. Fold changes and p-values for each of the 31 differential proteins are depicted in Table S2.

**Table 1:**
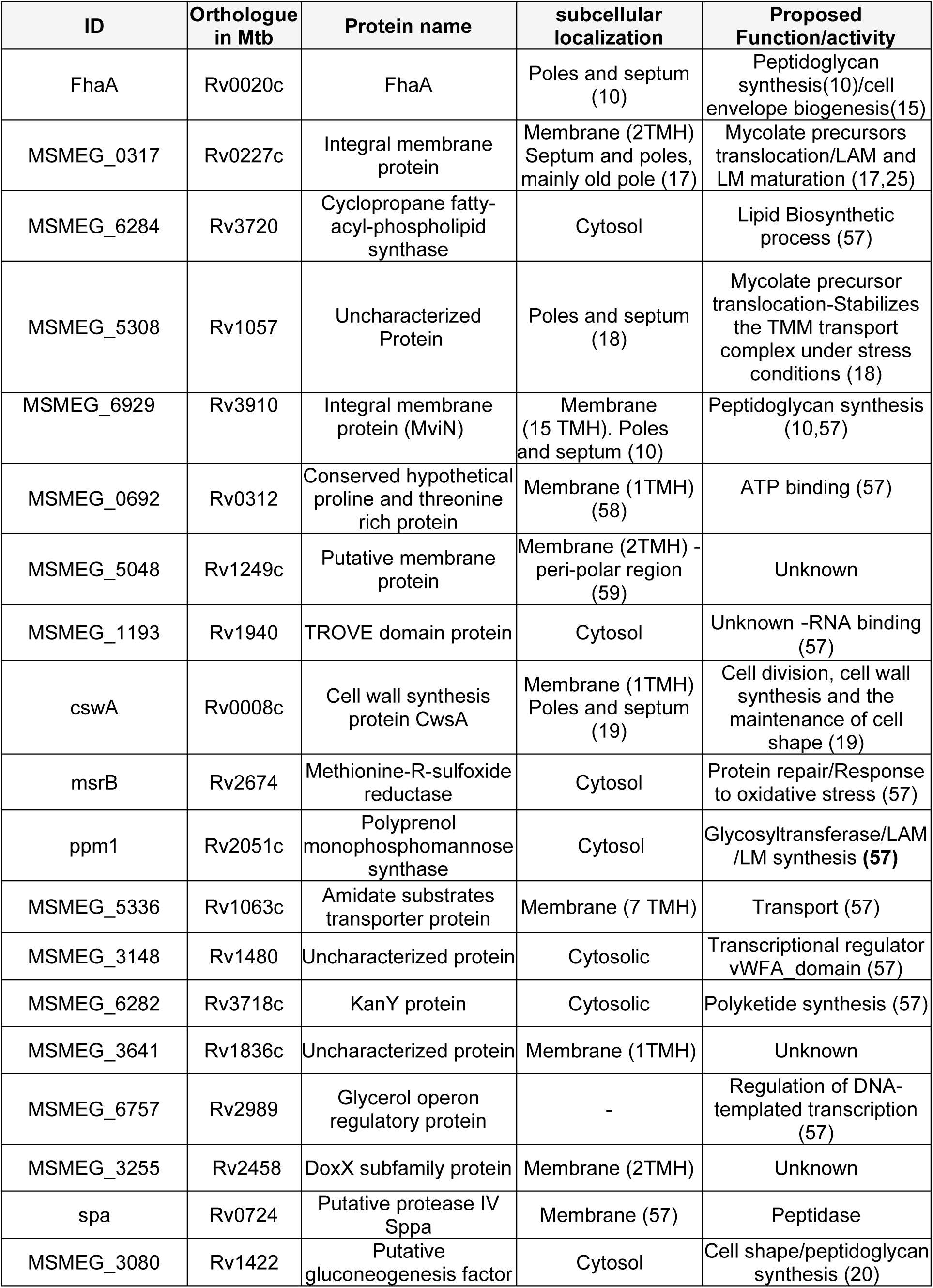
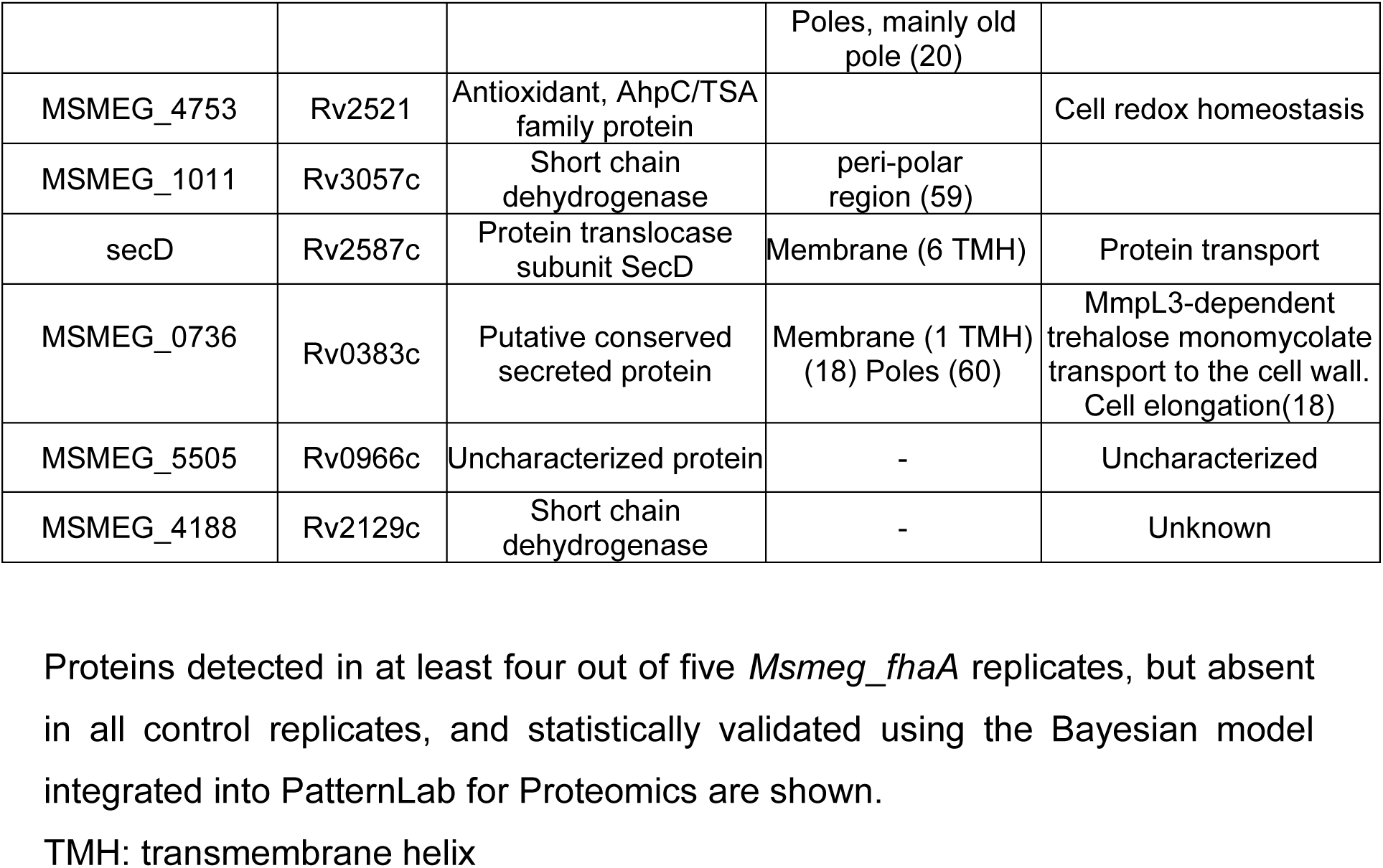
FhaA interactome: Proteins exclusively detected in FhaA mediated complexes.

| ID | Orthologue in Mtb | Protein name | subcellular localization | Proposed Function/activity |
| --- | --- | --- | --- | --- |
| FhaA | Rv0020c | FhaA | Poles and septum (10) | Peptidoglycan synthesis(10)/cell envelope biogenesis(15) |
| MSMEG_0317 | Rv0227c | Integral membrane protein | Membrane (2TMH)<br>Septum and poles, mainly old pole (17) | Mycolate precursors translocation/LAM and LM maturation (17,25) |
| MSMEG_6284 | Rv3720 | Cyclopropane fatty-acyl-phospholipid synthase | Cytosol | Lipid Biosynthetic process (57) |
| MSMEG_5308 | Rv1057 | Uncharacterized Protein | Poles and septum (18) | Mycolate precursor translocation-Stabilizes the TMM transport complex under stress conditions (18) |
| MSMEG_6929 | Rv3910 | Integral membrane protein (MviN) | Membrane (15 TMH). Poles and septum (10) | Peptidoglycan synthesis (10,57) |
| MSMEG_0692 | Rv0312 | Conserved hypothetical proline and threonine rich protein | Membrane (1TMH) (58) | ATP binding (57) |
| MSMEG_5048 | Rv1249c | Putative membrane protein | Membrane (2TMH) - peri-polar region (59) | Unknown |
| MSMEG_1193 | Rv1940 | TROVE domain protein | Cytosol | Unknown -RNA binding (57) |
| cswA | Rv0008c | Cell wall synthesis protein CswA | Membrane (1TMH)<br>Poles and septum (19) | Cell division, cell wall synthesis and the maintenance of cell shape (19) |
| msrB | Rv2674 | Methionine-R-sulfoxide reductase | Cytosol | Protein repair/Response to oxidative stress (57) |
| ppm1 | Rv2051c | Polyprenol monophosphomannose synthase | Cytosol | Glycosyltransferase/LAM /LM synthesis (57) |
| MSMEG_5336 | Rv1063c | Amidate substrates transporter protein | Membrane (7 TMH) | Transport (57) |
| MSMEG_3148 | Rv1480 | Uncharacterized protein | Cytosolic | Transcriptional regulator vWFA_domain (57) |
| MSMEG_6282 | Rv3718c | KanY protein | Cytosolic | Polyketide synthesis (57) |
| MSMEG_3641 | Rv1836c | Uncharacterized protein | Membrane (1TMH) | Unknown |
| MSMEG_6757 | Rv2989 | Glycerol operon regulatory protein | - | Regulation of DNA-templated transcription (57) |
| MSMEG_3255 | Rv2458 | DoxX subfamily protein | Membrane (2TMH) | Unknown |
| spa | Rv0724 | Putative protease IV Sppa | Membrane (57) | Peptidase |
| MSMEG_3080 | Rv1422 | Putative gluconeogenesis factor | Cytosol | Cell shape/peptidoglycan synthesis (20) |
|  |  |  | Poles, mainly old pole (20) |  |
| MSMEG_4753 | Rv2521 | Antioxidant, AhpC/TSA family protein |  | Cell redox homeostasis |
| MSMEG_1011 | Rv3057c | Short chain dehydrogenase | peri-polar region (59) |  |
| secD | Rv2587c | Protein translocase subunit SecD | Membrane (6 TMH) | Protein transport |
| MSMEG_0736 | Rv0383c | Putative conserved secreted protein | Membrane (1 TMH) (18) Poles (60) | MmpL3-dependent trehalose monomycolate transport to the cell wall. Cell elongation(18) |
| MSMEG_5505 | Rv0966c | Uncharacterized protein | - | Uncharacterized |
| MSMEG_4188 | Rv2129c | Short chain dehydrogenase | - | Unknown |

Proteins detected in at least four out of five *Msmeg\_fhaA* replicates, but absent in all control replicates, and statistically validated using the Bayesian model integrated into PatternLab for Proteomics are shown.
TMH: transmembrane helix

### Proteins recovered from FhaA interactome are related to cell division/elongation and cell envelope biogenesis

A detailed analysis of the FhaA interactors sheds light on its possible functions. The most abundant protein in the FhaA interactome, MSMEG_0317, is the integral membrane protein PgfA (for Polar Growth Factor A) that was recently identified as being crucial for growth from the old pole (17). PfgA also interacts with MmpL3, the trehalose monomycolate (TMM) transporter that plays an important role in mycolic acid trafficking across the membrane and cell envelope composition (17,18). Interestingly, two additional FhaA interactors participate in the regulation of MmpL3-mediated mycolic acid translocation: MSMEG_5308 and MSMEG_0736, with the latter being renamed as TtfA (for TMM transport factor A) (18). Overall, the interactome includes 9 previously reported MmpL3 interactors (18,21) in addition to MmpL3 itself. Proteins that participate in the biosynthesis of the different layers of the complex mycobacterial cell wall were also recovered as FhaA direct/indirect interactors.

In addition to Mvin (10), the list includes CwsA (for Cell Wall Synthesis protein A) (22), proteins that participate in lipomannan (LM) and lipoarabinomannan (LAM) biosynthesis such as the polyprenyl monophosphomannose synthase Ppm1 (23,24) and MSMEG_0317 (17,25), or yet the transcriptional regulator WhiA and the DivIVA domain-containing protein SepIVA (MSMEG_2416), both involved in cell division, cell length and/or cell envelope biosynthesis (26,27). Finally, the interactome also includes the scaffolds of the divisome and elongasome machineries (FtsZ and Wag31 respectively). Taken together, these results strongly support the involvement of FhaA in mycobacterial cell envelope biosynthesis during cell growth and division.

### FhaA overexpression alters cell envelope composition/structure

The overexpression of FhaA leads to a significant decrease in cell surface hydrophobicity (Figure 2A), supporting the hypothesis that this protein is involved in cell wall biogenesis. As this is a physicochemical property pivotal for cell-cell and cell-surface adhesion behaviours (28,29), we investigated the impact of FhaA overexpression on the formation of multicellular structures and observed that the *Msmeg_fhaA* strain has an impaired capability of biofilm formation (Figure 2B), which is not related to defects in the final biomass reached (Figure S1).

**Figure 2.**
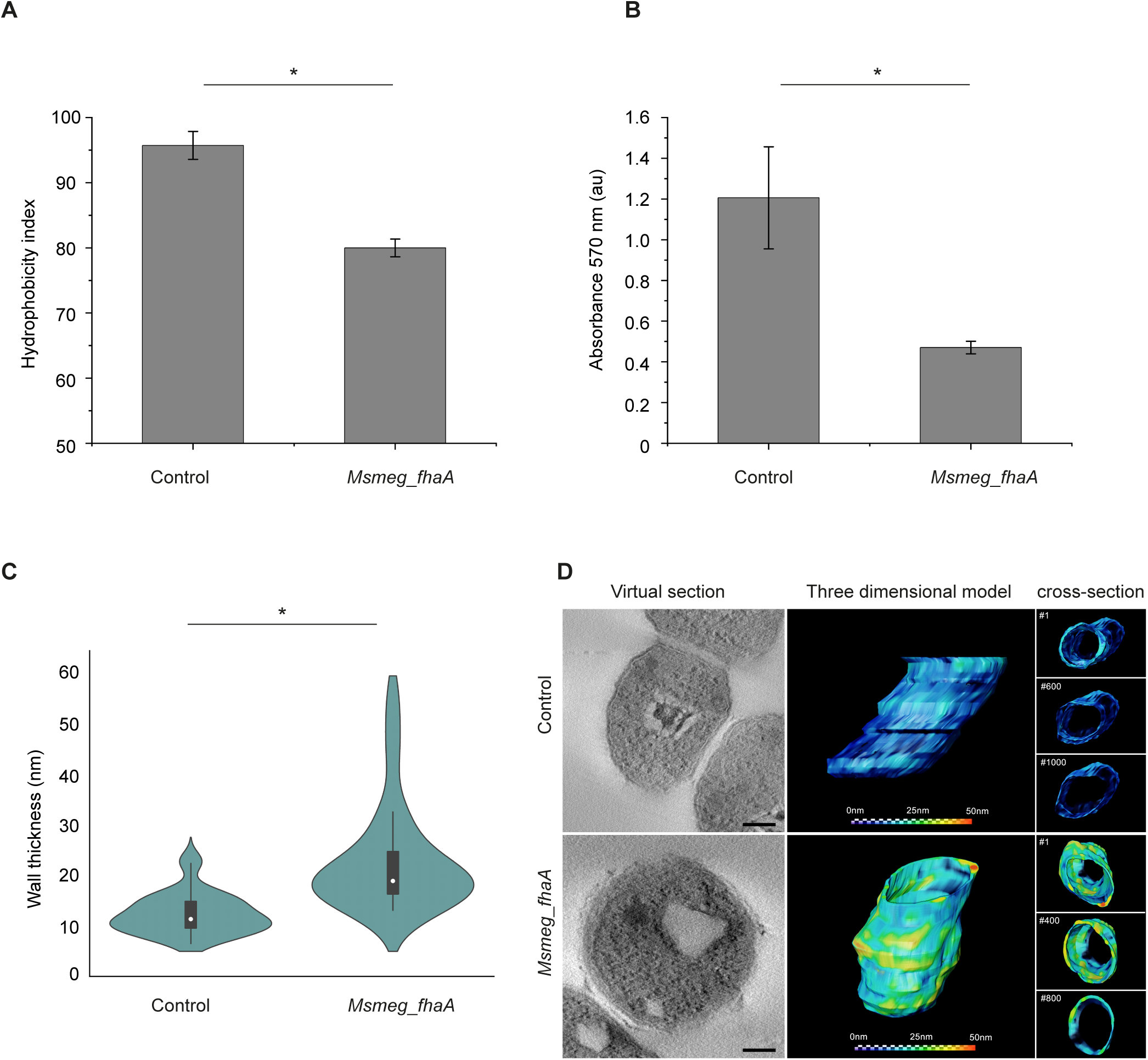
FhaA overexpression alters mycobacteria cell surface and cell envelope composition/structure. **A)** Cell surface hydrophobicity test. The figure shows partitioning of control and *Msmeg*_*fhaA* strains between PBS and xylene. The graph depicts the hydrophobicity index, defined as the percentage of the initial aqueous layer absorbance retained in the xylene fraction after partitioning. Assays were performed by triplicate Mean ± SD; *Indicates statistically significant difference determined by ANOVA, p < 0.05. **B)** Biofilm formation assay. Biofilm formation was evaluated in 96 well plates, by staining biofilms with crystal violet and measuring absorbance at 570 nm. Mean ± SD; *Indicates statistically significant difference determined by ANOVA, p < 0.05. **C)** Graphic comparison of the average cell wall thickness measured from TEM images (Figure S 2) for each strain. Violin plot illustrates the distribution of wall thickness, supporting the heterogeneity observed in the *Msmeg*_*fhaA* strain. Kolmogorov-Smirnov was applied; n = 30 cells for each group; * = p<0.05. White circles represent median; grey boxes represent 25%-75% percentile; values outside whiskers represent outliers. **D)** Thickness map representing variations in cell wall thickness across the cell volume. The color intensity corresponds to the magnitude of thickness, where warmer hues indicate greater thickness (25 – 50 nm) and cooler hues denote thinner regions (0 -25 nm). Left panel: virtual sections from representative control and *Msmeg*_*fhaA* tomograms utilized in the modeling of the cell walls. Right panel: Three-dimensional model of partial volumes of control and *Msmeg*_*fhaA* cells. A predominant dark blue phenotype throughout the volume is observed for control strain while there is a prevalence of warm hues along the majority of the *Msmeg*_*fhaA* sampled volume, indicating the increase in the wall thicknesses. On the right side, cross section view of different sequential slices along the Z axis. Numbers indicate where the models were sectioned. Scale bar: 100 nm.

Transmission electron microscopy (TEM) images confirmed that the strain overexpressing FhaA exhibits an abnormal cell envelope (Figure 2C and Figure S2). The images show areas that have an unusually thick cell wall that appear as electron lucid blobs with aberrant distribution (Figure S2). Compared to the control, there is an increase in the average thickness of the cell envelope (Figure 2C-D), with these thickened areas being heterogeneously distributed across the surface. (Figure S2). Cell wall maps across the cell volume obtained by electron tomography (ET) further confirm the alterations in the *Msmeg*_*fhaA* cell envelope and highlight the thickness heterogeneity along the cell volume (Figure 2D).

Further evidence for the effect of FhaA overexpression on cell envelope comes from the analysis of membrane properties of the two strains. We used scanning confocal microscopy to image cells previously treated with LAURDAN, an amphiphilic fluorescent dye that penetrates the membrane lipids and whose emission spectra change according to environment molecular composition and polarity (30,31). When plotted on a diagram, phasors corresponding to the control strain tend to cluster at higher angles and closer to the origin of the axes compared to those representing *Msmeg_fhaA* (Figure 3). In addition, there is a change on the profile on the linear combination obtained at the phasor plot for the two strains. Thus, the misalignment between the two trajectories plus the spectral shift and broadening observed for the strain overexpressing FhaA can be attributed to changes in the molecular environment sensed by LAURDAN (Figure 3B).

**Figure 3.**
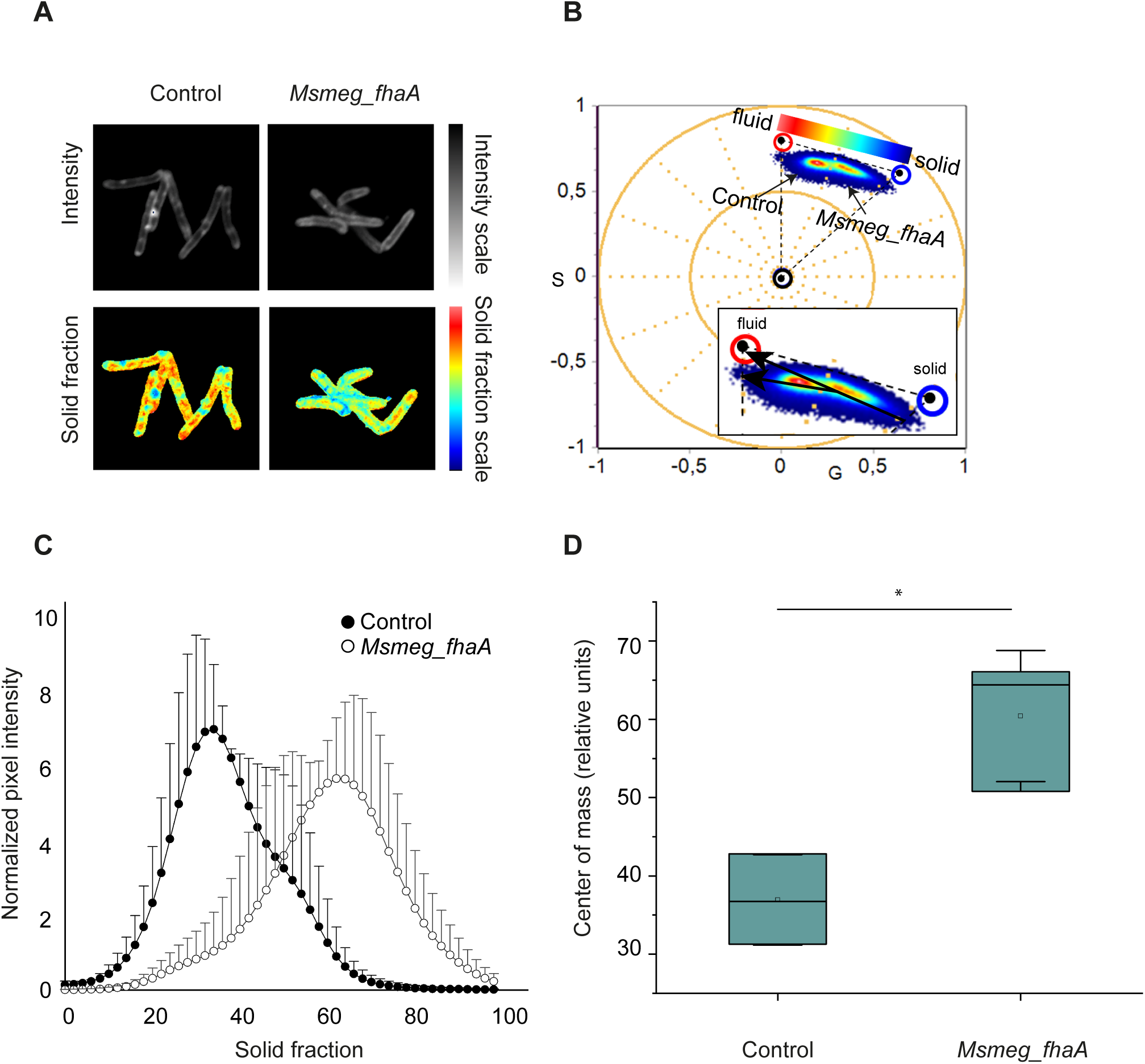
Scanning confocal microscopy using the fluorescent dye LAURDAN. **A)** Representative images for intensity and pseudocolor image of LAURDAN from of control and Msmeg_fhaA strains. Pseudocolor images were generated by using the color scale indicated on B and represents spectral shift from blue to red. **B)** Spectral phasor plot of LAURDAN fluorescence emission from control and *Msmeg*_*fhaA* strains. Emission spectra were Fourier transformed into the G and S (corresponding to the real and imaginary parts of the first harmonic of the Fourier transform) to obtain the spectral phasor plot. Data indicates strong differences in envelope fluidity between strains, as measured by LAURDAN emission. The *Msmeg*_*fhaA* strain clusters are shifted clockwise (blue-shifted) relative to the control strain and are further from the plot origin (indicating spectral widening). Additionally, the amplified section shows two different trajectories corresponding to each strain, clearly indicating different molecular environments for LAURDAN. **C)** Plots illustrating normalized pixel intensity vs. solid fraction. Black dots represent the control strain; white dots represent the *Msmeg_fhaA* strain. **D)** Box plot representing the values of the center of mass for the curves depicted in C.

Altogether our results indicate that FhaA overexpression has important impacts on mycobacterial cell envelope composition and/or structure.

### The overexpression of FhaA affects cell morphology

To investigate the effect of FhaA on elongation we evaluated cell morphology of *Msmeg_fhaA.* Confocal microscopy analysis of bacteria stained with Sulforhodamine-DHPE revealed that the overexpression of FhaA led to significantly shorter cells, exhibiting an average length of 4.5 ± 0.1µm (average length of control: 7.0 ± 0.2 µm) (Figure 4A). This observation was subsequently corroborated through scanning electron microscopy (SEM), which revealed that *Msmeg_fhaA* cells exhibit abnormal and heterogeneous morphology and dimensions, marked by shorter and wider cells with defects at poles and septum, the places where new cell wall material is incorporated. While in some cases swelling at the septum was observed, the vast majority of the cells presented defects at the poles (Figure 4B-C). The aberrant morphology is distinguished by the thickening and curling of bacterial poles, with swollen and bulged tips that suggest compromised polar cell envelope integrity. Interestingly, these defects were mainly asymmetrical, being observed at one of the cell poles, with fewer cells presenting alterations at both poles (Figure 4B). Ultrastructural analysis of cell poles by ET showed three cell wall layers as expected, with the middle layer displaying increased thickness in *Msmeg_fhaA* when compared to the control strain (Figure 4D, Supplementary video), suggesting an altered synthesis of the layers between the mycomembrane and inner membrane. The virtual section of a cell tomogram from an aberrant *Msmeg*_*fhaA* pole corroborated the thickening of the cell wall (Figure 4D-E). These observed morphological alterations thus suggest that FhaA overexpression disrupts normal polar cell elongation and cell wall synthesis.

**Figure 4.**
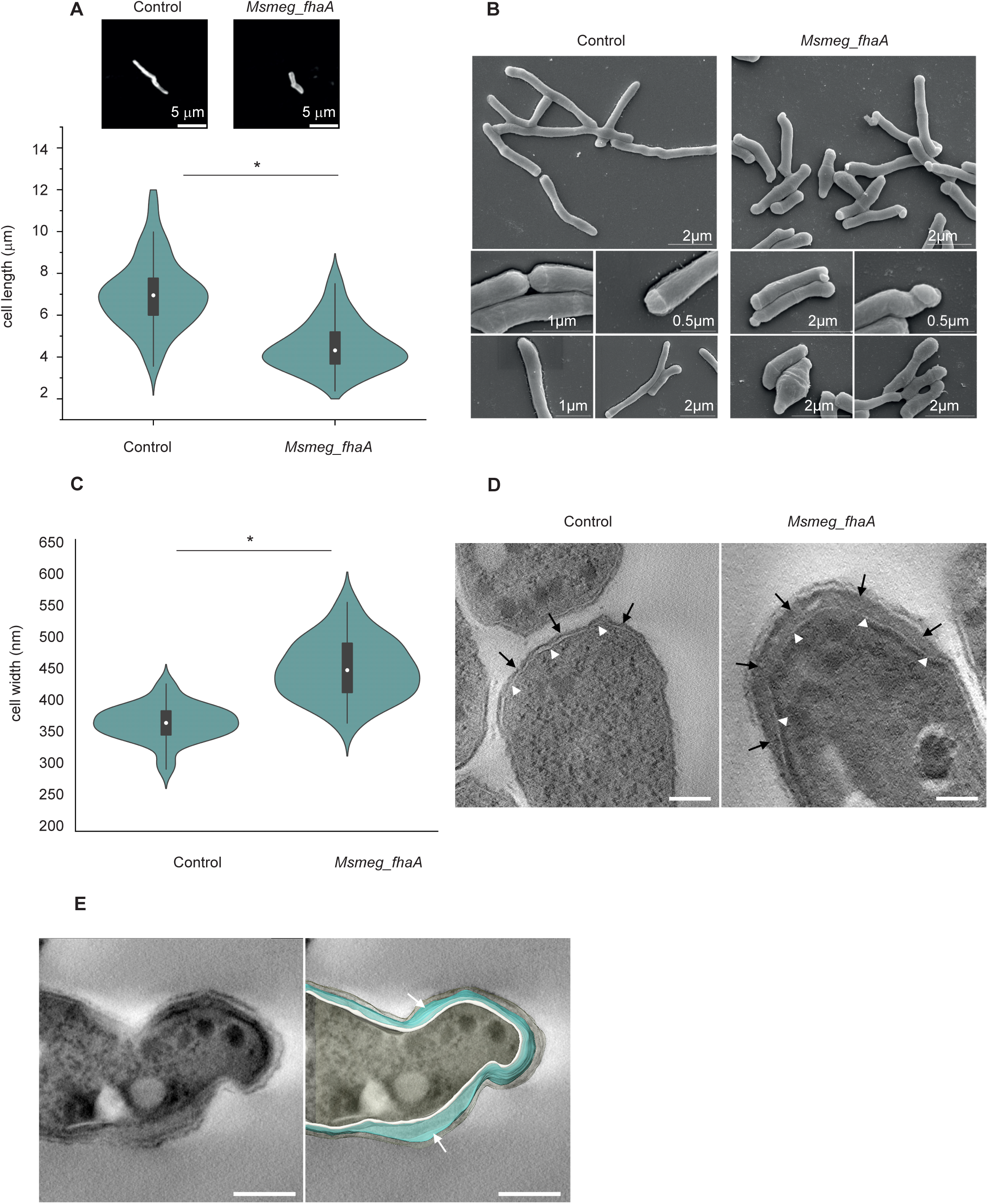
FhaA overexpression alters cell morphology. **A)** Violin plot and representative images of Sulforhodamine-DHPE stained bacteria illustrating differences in cell length between both strains. Average length is 7.0 ± 0.2 μm for control strain and 4.5 ± 0.1μm for *Msmeg_fhaA* strain. *Indicates statistically significant difference determined by Two samples T-test. P < 0.05; n>100 cells for each group. White circles represent median; grey boxes represent 25%-75% percentile; values outside whiskers represent outliers. **B)** Scanning electron microscopy showing morphological differences between strains. Images of *Msmeg*_*fhaA* strain reveal a heterogeneity in cell shapes, length, and width when compared to control. In addition, most of the *Msmeg*_*fhaA* cells exhibit one aberrant pole. **C)** Violin plot showing that cell width is altered in *Msmeg*_*fhaA* strain. Measurements of cell width were performed from SEM images. *Indicates statistically significant difference determined by one-way ANOVA. p < 0.05; n=30 cells for each group. White circles represent median; grey boxes represent 25%-75% percentile; values outside whiskers represent outliers. **D)** Virtual sections from tomograms of control and *Msmeg*_*fhaA* strains showing ultrastructural differences at cell tips. White arrowheads indicate the plasma membrane. Notably, an increase in the middle layer (black arrows) is observed within the cell wall of *Msmeg*_*fhaA*, contrasting with the consistently thinner layer exhibited in the control. Scale bar: 100 nm. **E)** Left: Virtual section of a *Msmeg*_*fhaA* cell tomogram with a ‘curved’ tip. Right: Top view of the 3D model, emphasizing the thickening of the cell wall (white arrows), which potentially alters the cell topography near the tip, contributing to the observed curved phenotype. White layer represents the plasma membrane; light blue indicates peptidoglycan/arabinogalactan; light yellow indicates outer membrane. Scale bar: 200 nm.

### FhaA overexpression leads to the mislocalization of PG biosynthesis

Next, we evaluated the effect of FhaA overexpression on PG synthesis using the fluorescent D-amino acid analogue HADA (32) to label the nascent PG. As expected, new cell wall material in the control strain is specifically inserted at the poles and septum (Figure 5A). However, in the *Msmeg_fhaA* strain, PG synthesis is not strictly confined to these sites, as HADA is additionally incorporated into discrete foci along the cell surface (Figure 5A). To quantify the extent of PG synthesis delocalization, we assessed the distance between focal points of HADA incorporation in each bacterium, normalized to the cell length. As expected, we detected 2 or 3 local maxima of fluorescence intensity for the control strain (Figure 5A and C) corresponding to the two poles (non-septate cells), or to the two poles plus the septum (septate cells), respectively (average number of HADA foci per cell: 2.7±0.5). In this case, the interspace between foci of PG synthesis correlates with the pole-septum or pole-pole distances as expected (Figure 5B-C). For *Msmeg_fhaA,* the average number of foci per cell increases to 4.5 ±1.7, and the relative distance between foci is significantly shorter, indicating that the PG biosynthetic machinery is mislocalized (Figure 5B-C).

**Figura 5.**
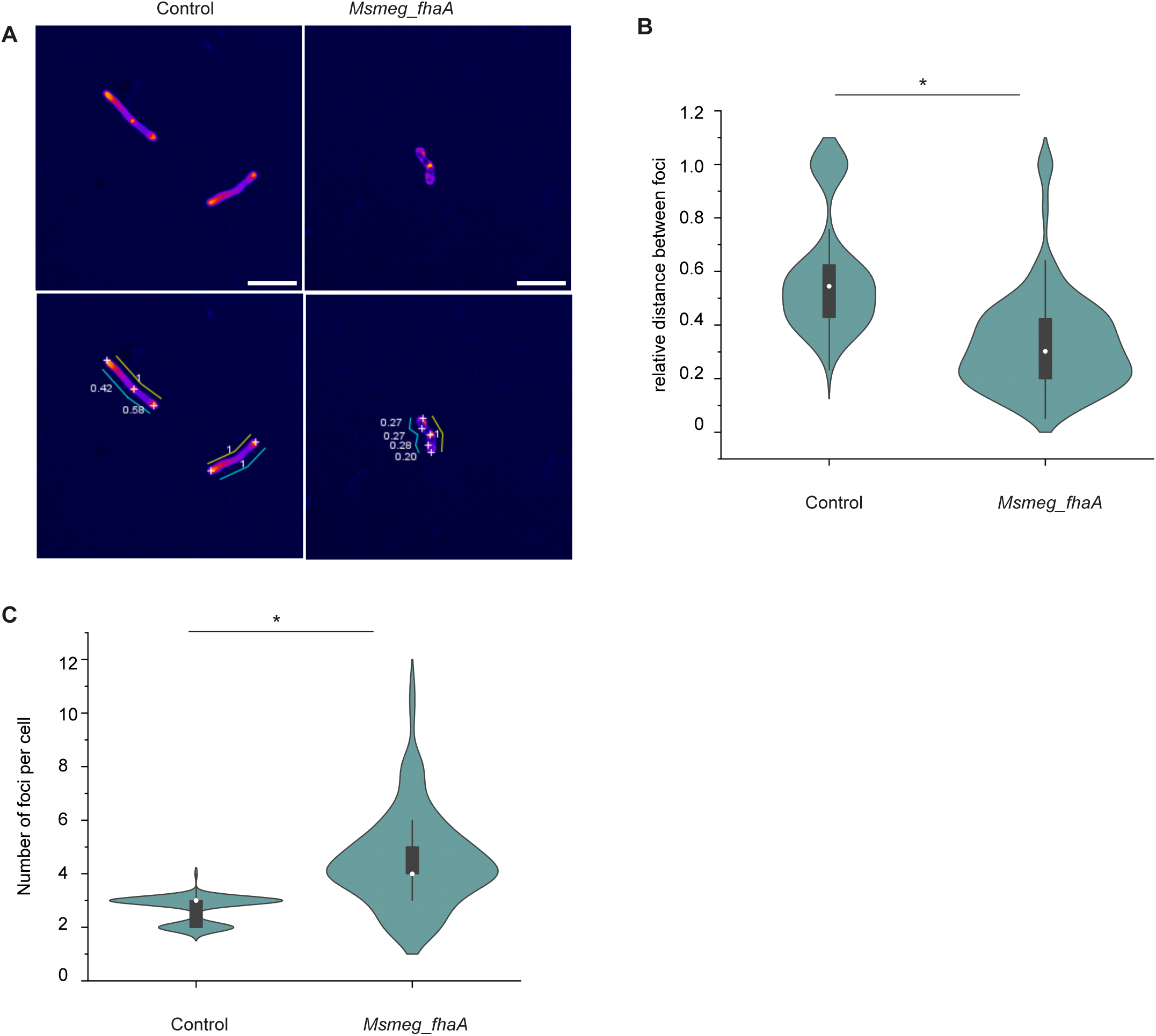
FhaA overexpression leads to mislocalization of the PG synthesis machinery. **A)** Representative images of control and *Msmeg*_*fhaA* strains showing PG synthesis distribution. Fire LUT was applied to HADA signal to enhance visibility of regions with higher fluorescence intensity. *Find maxima* tool of Image J was used to detect local intensity maxima for the HADA signal (white crosses) and distances between focuses were measured (cyan sticks). As the cell length is significantly different among both strains, distances between foci were relativized to cell length (yellow sticks). Scale bars: 5 µm. **B)** Violin plot showing the differences in distribution of distances between foci for control and *Msmeg*_*fhaA* strain. Distance between foci (poles and septa) for septate control strain oscillates between 50/50 and 70/30 of the total cell length, as expected. *Indicates statistically significant difference determined by Kolmogorov-Smirnov. p < 0.05; n>100 cells for each group. White circles represent median; grey boxes represent 25%-75% percentile; values outside whiskers represent outliers. **C)** Violin plot showing that number of HADA foci per cell is increased in *Msmeg*_*fhaA* strain. Control cells exhibit 2 foci (both poles, non-septate bacteria) or three, (two poles and septum, septate bacteria), *Msmeg*_*fhaA* cells exhibit multiple foci, even when non septate. *Indicates statistically significant difference determined by Kolmogorov-Smirnov, p < 0.05; n >100 cells for each group. White circles represent median; grey boxes represent 25%-75% percentile; values outside whiskers represent outliers.

The abnormal localization of the cell wall synthesis machinery, leading to bulges and branches, was previously shown for *M. smegmatis* strains overexpressing the elongasome scaffold Wag31 (33). To evaluate if the delocalization of the PG synthesis machinery could be associated with increased levels of Wag 31 in *Msmeg*_*fhaA*, we performed a comparative analysis by mass spectrometry. The results confirmed the overexpression of FhaA, but Wag31 levels were not statistically different between *Msmeg*_*fhaA* and control strain (Table S4). This result suggests that the elevated levels of FhaA could be the primary factor driving the delocalization of cell wall biosynthesis machinery.

### FhaA is necessary for asymmetric polar elongation

To further investigate the role of FhaA in elongation, we evaluated the cell morphology and HADA incorporation for a strain lacking FhaA *(Msmeg_*Δ*fhaA*) In accordance with a previous report (15), *Msmeg_ΔfhaA* cells were shorter than the WT strain, and cell length was partially recovered after complementation (Figure 6A) confirming a role for FhaA in cell elongation.

**Figure 6:**
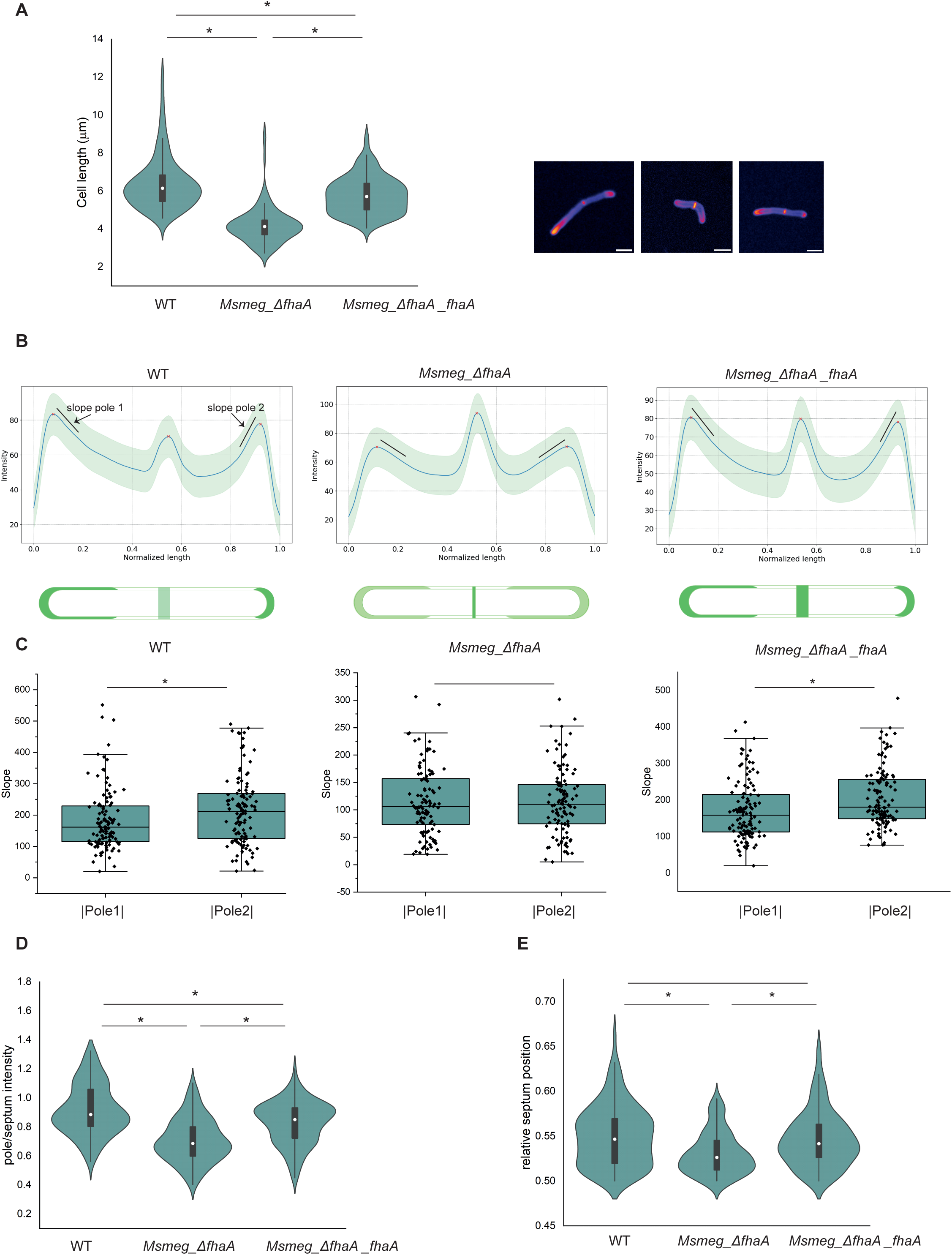
FhaA is necessary for asymmetric polar elongation. **A).** Violin plot and representative images showing differences in length between strains. Msmeg_ΔfhaA cells are shorter than WT cells and length is partially recovered after complementation. * Indicates statistically significant difference determined by Kruskal-Wallis test, p < 0.05; n >100 cells for each group. White circles represent median; grey boxes represent 25%-75% percentile; values outside whiskers represents outliers. Scale bar 2 μm **B**). Average HADA fluorescence profiles along the cell for >100 septate cells. Blurred zone represents standard deviation. Profiles consist in 3 peaks corresponding to both poles, (named pole 1 and pole 2) and septum. For WT maximum intensity is located at poles, while for Msmeg_ΔfhaA, strain the maximum of intensity is located at septum. *Msmeg_ΔfhaA_fhaA* exhibits an intermediate phenotype exhibiting 3 peaks of comparable intensity. Schemes below represents HADA deposition patterns for each strain. **C)** Box plots showing the slope (black lines in fluorescence profiles) at pole 1 and pole 2 for >100 cells allowed to corroborate the asymmetric growth for WT and *Msmeg_ΔfhaA_fhaA*. For *Msmeg_ΔfhaA*. HADA incorporation at both poles was undistinguishable. * Indicates statistically significant difference determined by Kolmogorov-Smirnov test, p < 0.05; n >100 cells for each group. Box represents 25%-75% percentile and median; values outside whiskers represents outliers. **D**). Violin plot illustrating the ratio between intensity at poles (average of both) and intensity at septum. *Indicates statistically significant difference determined by one-way ANOVA. p < 0.05; n>100 cells for each group. White circles represent median; grey boxes represent 25%-75% percentile; values outside whiskers represent outliers. E). Violin plot showing the distribution of the relative septum positionin WT, *Msmeg_ΔfhaA* and *Msmeg_ΔfhaA_fhaA* strains. The asymmetrical position of the septum is lost in *Msmeg_ΔfhaA* strain, and it is completely restored after complementation. *Indicates statistically significant difference determined by one-way ANOVA. p < 0.05; n>100 cells for each group. White circles represent median; grey boxes represent 25%-75% percentile; values outside whiskers represent outliers.

Fluorescence intensity profiles in dividing WT cells revealed the presence of three maxima at septum and poles (Figure 6B), with the poles exhibiting a greater intensity and asymmetric elongation, as reported previously (6,34,35). The faster growing pole showed HADA signals covering a broader area from the tip, and a smaller slope of fluorescence intensity (Figure 6A and 6B). Conversely, the fluorescence signal at the slower growing pole appears concentrated within a more restricted region and the slope in fluorescence intensity profile is higher (Figure 6A-C), confirming a statistically significant difference in the extent of HADA incorporation for the WT strain (Figure 6A and 6C). However, *Msmeg_ΔfhaA* displays a distinct HADA profile characterized by an increased intensity at the septum and significantly lower levels of HADA incorporation at the poles (Figure 6B). The ratio of pole intensity/septum intensity (with pole intensity as the average of both poles) is significantly different between WT and *Msmeg_ΔfhaA*, and this abnormal distribution of PG synthesis is partially reverted after complementation (Figure 6D, Figure S3 A). In addition, the polar incorporation of HADA by *Msmeg_ΔfhaA* encompasses a broader area at both poles, when compared with WT. Surprisingly, the deletion of FhaA not only altered the pattern and quantity of HADA incorporation but also resulted in a symmetrical polar incorporation of new cell wall material, as revealed by the average fluorescence profile and the similar slope of fluorescence intensity at each pole in *Msmeg_*Δ*fhaA* (Figure 6B and 6C). Moreover, this slope is smaller than that measured for the fast-growing pole of WT which, together with the observation that *Msmeg_*Δ*fhaA* cells are shorter, indicates a more diffuse localization of the PG synthesis machinery (Figure 6B-C and S3 B). It is important to note that complementation of *Msmeg_ΔfhaA* completely recovers polar cell wall synthesis and asymmetric PG incorporation (Figure 6B-C and Figure S3 A-B).

As a consequence of the well-documented differences in growth rates between the old and new poles, mycobacteria exhibit an asymmetric position of the septum and considerable variability in cell size among the population. Thus, we assessed septum position and cell length variability in *Msmeg*_Δ*fhaA.* Consistent with the loss of asymmetric growth, in the *fhaA* deletion strain the septum is positioned closer to the midcell compared to WT, while the septal position asymmetry is restored after complementation (Figure 6E). The loss of asymmetric growth in *Msmeg_*Δ*fhaA* is further confirmed by a more homogeneous population in length, compared to either the WT or the complemented strain (Figure 6A). Moreover, using a *M. smegmatis* strain overexpressing FhaA fused to the fluorescent protein mScarlet (*Msmeg*_*mscarlet*_*fhaA*), we showed that the protein localizes to the poles and septa as anticipated, with a predominant localization at the old pole that matches the higher HADA incorporation pattern at this site (Figure 7).

**Figure 7:**
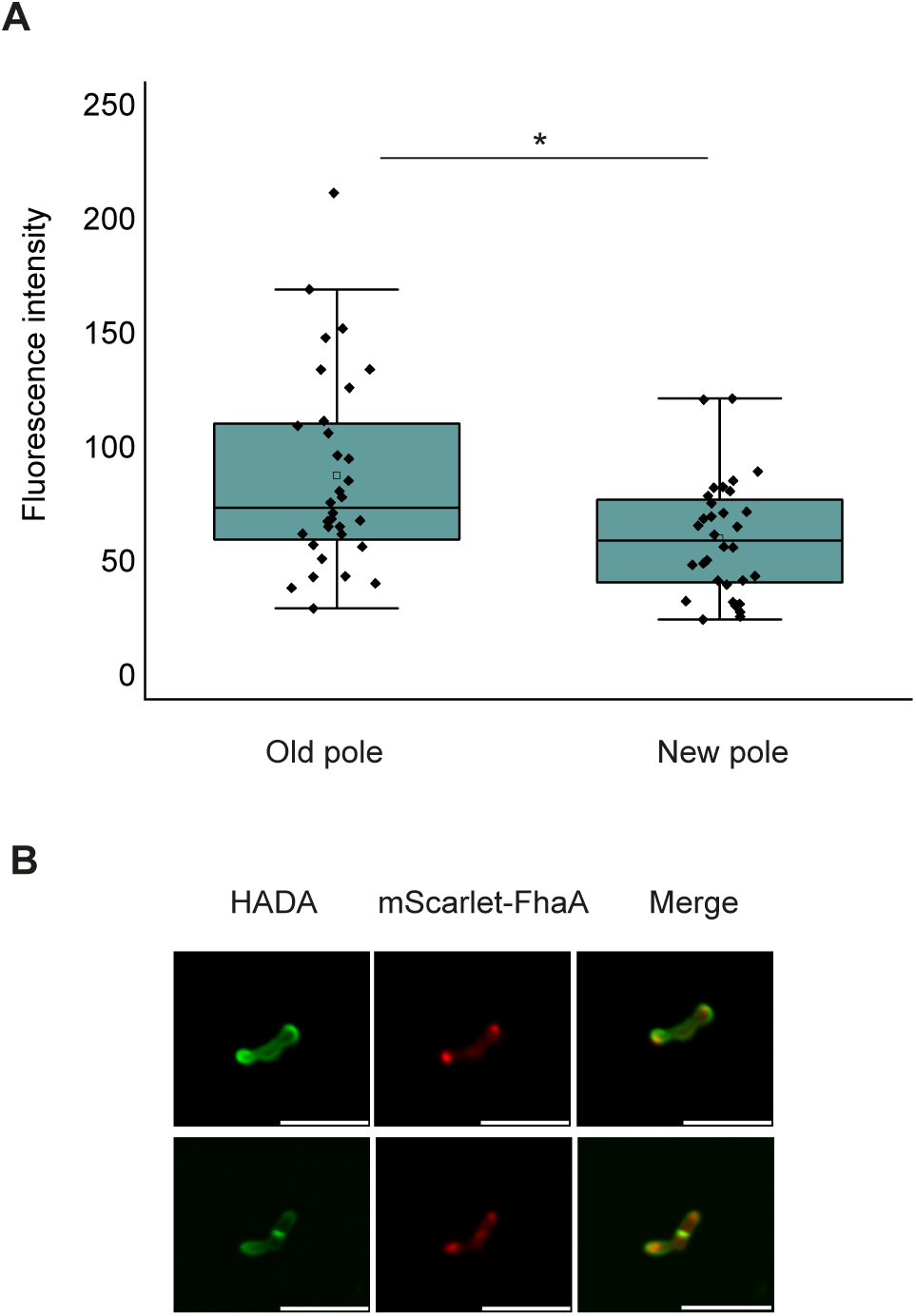
FhaA localizes preferentially at the old pole. **A)**. Box plot showing differences in distribution of mScarlet-FhaA fluorescence intensity at both poles. Poles were classified as old and new based on the pattern of HADA fluorescence incorporation. The top asterisk indicates statistically significant difference determined by two samples T - test,<0.05; n=30 cells. White circles represent median; grey boxes represent 25%-75% percentile; values outside whiskers represent outliers. **B**). Representative images of an *Msmeg_ mscarlet_fhaA* cell showing colocalization of HADA signal and mScarlet-FhaA at poles and septum. mScarlet-FhaA accumulates preferentially at the old pole. Scale bar: 5μm.

## Discussion

The cellular growth of mycobacteria is distinguished by the incorporation of new cell wall material at the poles in an asymmetric manner, with the old pole inherited from the mother cell growing faster than the new pole formed after the last division. This asymmetry leads to a heterogeneous population in terms of size, growth rate and antibiotic susceptibility, which is critical for *M. tuberculosis* pathogenesis and the development of antibiotic-resistant strains. Despite its importance, the molecular mechanisms that sustain asymmetric polar growth are still not well understood. Here, we present strong evidence that FhaA is a key elongation factor predominantly localizing to the old pole, and that is crucial for cell envelope integrity and the asymmetry of polar growth.

The FhaA architecture of two modular domains separated by a flexible linker is highly reminiscent of eukaryotic scaffolding proteins (10,13). This protein organization suggests that FhaA could have a tethering role, bringing together different molecular machineries. Our interactomic analysis shows that FhaA is part of a protein network involved in the biosynthesis of different layers of the complex mycobacterial cell envelope, including proteins associated with PG, LM/LAM, and mycolic acid synthesis and transport. These findings are in line with, and extend, the previous observation that FhaA interacts with the Lipid II flippase Mvin (10). Consistently, the FhaA interactome significantly overlaps with those reported for other well-known components of the *Mycobacteriales* elongasome/divisome, namely MmpL3 and FtsQ (18,21,36) (Tables S1 and S2), and several of the proteins identified in our work (FhaA itself, MSMEG_0317, MSMEG_ 5048, atpC, MmpL3, NusA and WhiA) were shown to interact with mycolates *in vivo* using photoactivatable TMM analogues (37) (Tables S1 and S2).

In line with these results, overexpression of FhaA led to alterations in the cell envelope composition and structure. The decreased surface hydrophobicity, along with the impairment in biofilm formation of the *Msmeg*_*fhaA* strain, points to defective biosynthesis or stability of the mycomembrane and/or the underlying layers of the cell wall. Release of significant amounts of membrane fragments containing mycolic acid esters of trehalose, as a result of impaired mycomembrane stability has been reported for a *Corynebacterium glutamicum* strain defective in arabinogalactan synthesis (38). Our ET analysis reveals irregular thickening of the cell wall layer between the two membranes in the strain overexpressing FhaA, providing a potential explanation for the observed phenotypes. Finally, our analysis of LAURDAN fluorescence indicates an altered fluidity of the cell envelope lipidic layers. Nevertheless, considering our data as a whole, we are inclined to speculate that the perturbation of the intermediate layers of the cell envelope could be responsible for the change in polarity and water relaxation sensed by LAURDAN. Altogether, our interactome and phenotypic characterization strongly indicates that FhaA is a part of the molecular machinery responsible for the synthesis of the complex cell envelope of mycobacteria and plays a functional role in this biosynthetic process.

We confirmed that FhaA localizes preferentially to the poles and septum, as previously reported (10). Furthermore, our quantitative analyses revealed an asymmetric distribution between the poles, with a higher concentration of FhaA specifically at the old pole. During growth and division PG synthesis is orchestrated by two multiprotein complexes: the elongasome responsible for polar elongation and the divisome responsible for cell division and septation. The short cell phenotype of *Msmeg*_Δ*fhaA* cells, together with a lower HADA incorporation at the poles, clearly indicates that FhaA partakes in polar PG synthesis and normal cell elongation. A previous report showed that FhaA depletion results in increased accumulation of nascent PG stem peptides at the cell poles and septum and thus propose that FhaA inhibits the late stages of PG biosynthesis via its interaction with Mvin (10). Our results consistently support a role for FhaA in this process, but based on HADA incorporation and morphological characterizations, we hypothesize that FhaA promotes PG synthesis. In addition, the characteristic asymmetric growth pattern of mycobacteria is lost in the absence of FhaA. Collectively, our results establish FhaA as a bona fide functional partner of the elongasome, essential for asymmetric polar elongation. Further supporting this hypothesis, the recruitment to the old pole of the FhaA’s top interactor PgfA, which shares the same localization pattern as FhaA, is known to be essential for establishing cellular asymmetry (17). The uneven distribution of key components of the cell elongation machinery, predominantly concentrated at the old pole, has been previously demonstrated and provides a plausible explanation for the differential polar growth rates (5,39–42). A biphasic growth model has been proposed, in which the new pole undergoes an initial phase of slower growth, during which Wag31 accumulates, followed by a period of rapid growth prior to the next division cycle (35). Another report suggests that the molecular basis for the polar growth of fast and slow poles are fundamentally different (17).

Various pieces of evidence in this work indicate that FhaA affects the precise subcellular localization of new cell wall material insertion. First, overexpression of FhaA leads to the insertion of HADA at multiple focal points along the cell length, as well as at the poles and septa, a phenotype that is not due to increased levels of Wag31 but seems to be directly linked to altered levels of FhaA. In parallel, TEM and ET reveal a heterogeneous cell wall with localized thickening, strongly suggesting that these enlarged areas of the cell wall can be correlated with the extra foci of HADA incorporation and the mislocalization of the elongasome machinery. Consistently, in the strain lacking FhaA, HADA incorporation extends over a broader region at the poles, surpassing even the area observed in the fast-growing pole of the WT strain. As *Msmeg*_*ΔfhaA* cells are shorter, the extended area of HADA incorporation suggests that the biosynthetic machinery is less positionally constrained at the poles, rather than indicating rapid growth. Taken together, our results indicate that FhaA participates in the regulation of the accurate localization of the elongasome and its biosynthetic activity.

Asymmetric growth is a key trait for mycobacteria adaptation and successful survival strategies, promoting heterogeneous populations with varied responses to environmental challenges and drugs. Thus, uncovering central molecular actors in this essential process deepens our understanding of mycobacterial biology while also identifying promising drug target candidates.

## MATERIALS AND METHODS

### Bacterial strains and growth conditions

The *M. smegmatis* overexpressing Strep-tagged FhaA (hereinafter referred to as *Msmeg*_*fhaA*), mScarlet-FhaA (referred as *Msmeg_mscarlet*_*fhaA*), and the control strain were obtained as previously described (43). Briefly, electrocompetent *Mycobacterium smegmatis* mc^2^ 155 were transformed with a pLAM12 plasmid containing the coding region of the gene Rv0020c (*fhaA* of *Mycobacterium tuberculosis*) plus an N-terminal tag (Strep-tag® II), or the gene Rv0020c plus the sequence of the red fluorescent protein mScarlet (44) at the N-terminus, both under the control of the *M. smegmatis* acetamidase promoter. As a control, *M. smegmatis* mc^2^ 155 transformed with an empty version of the pLAM12 plasmid was used (control). *M. smegmatis* strains were maintained on Middlebrook 7H10 agar plates (Difco) plus 10% ADC (0.2% dextrose, 0.5% bovine serum albumin, 0.085% NaCl). Liquid cultures were grown in Middlebrook 7H9 (Difco) plus 10% ADC and 0.05% (v/v) Tween 80® at 37°C and 220 r.p.m. until reaching an OD600 =0.8. All media were supplemented with kanamycin (50 μg/ml) and ampicillin (100 μg/ml). Expression of Strep-tag® II-FhaA was induced by addition of 0.2% acetamide during exponential growth phase (OD600 = 0.2). For interactomic analyses, five independent cultures of each strain were prepared. Strains of *M. smegmatis* mc^2^ 6 (WT strain in this work), mc^2^ 6_ΔfhaA (*Msmeg_ΔfhaA*), and mc^2^ 6_Δ*fhaA*_*pMV306_fhaA* (*Msmeg_ΔfhaA_fhaA*) were kindly provided by Dr. Raghunand Tirumalai (15). A table with all the strains used in this work is presented in Table S5.

### Chemical crosslinking in living cells

Chemical crosslinking was performed following the protocol previously used (45,46). Briefly, after 18h of induction, cultures were incubated with formaldehyde (final concentration 0.5%) at 37° C and 220 r.p.m. for 20 min, and the excess of formaldehyde was quenched by addition of 1/10 culture volume of ice-cold glycine (125 mM) in PBS for 20 min.

### Affinity purification of protein complexes

Cell cultures were harvested by centrifugation, washed with phosphate-buffered saline (PBS) and re-suspended in 25 mM HEPES, 150 mM NaCl, 1% glycerol, 1 mM EDTA, pH 7,4; 1x protease inhibitor cocktail (Roche®), 1 x phosphatase inhibitor (Sigma-Aldrich), 1.0% Triton X-100 (v/v). Lysates were obtained by sonication on ice (25% amplitude, 10 sec ON, 30 sec OFF. Total cycle: 8 min) followed by three cycles of 10 min vortexing in presence of glass beads (Glass beads, acid-washed ≤106 μm, Sigma-Aldrich) with 20 min intervals. After centrifugation, protein concentration of the supernatants was determined by densitometric analysis on SDS-PAGE gels.

FhaA protein complexes were purified using a commercial affinity resin (Strep-Tactin Sepharose®, IBA Solutions). Supplieŕs protocol was optimized, including extra washing steps with 1% Triton X-100 to decrease the number of unspecific interactions recovered. Elution was performed using D-desthiobiotin. As mock, protein extracts from control strain were submitted to the same purification protocol.

### Sample preparation for MS analysis

For interactome analysis purified complexes (5 replicates for each strain) were digested in solution at 37^0^C overnight, after Cys reduction (10 mM DTT) and alkylation (55 mM iodoacetamide). Peptide mixtures were desalted using micro-columns (C18 ZipTip®, Merck, Millipore), eluted in 0.1% formic acid in ACN, dried and resuspended in 0.1% formic acid prior to its analysis by nano-LC MS/MS.

### Nano-LC MS/MS analysis

Analysis of peptide mixtures was performed using liquid chromatography-tandem mass spectrometry (LC-MS/MS). Peptides were separated using a nano-HPLC system (EASY-nLC 1000, Thermo Scientific) equipped with a reverse-phase column (EASY-Spray column, 15 cm × 75 µm ID, PepMap RSLC C18, 2 µm, Thermo Scientific) and a precolumn (Acclaim Pepmap 100 C18 3 µm, 75 µm x 20 mm, Thermo). Elution was performed at a constant flow rate of 250 nL/min with a two-solvent system (A: 0.1% formic acid in water and B: 0.1% formic acid in acetonitrile) and the following gradient: 0% to 50% B over 100 min, 50% to 100% B over 10 min. Column temperature was set to 45 °C. For total proteome analysis, 3 replicates of each strain were used. 68 mg of total protein extracts were loaded on SDS PAGE (12%). The gel was fixed, stained and each lane was excised into 4 gel pieces that were processed for MS analysis as previously described (47). Briefly, Cys reduction and alkylation was performed by incubation with 10 mM DTT at 56 °C followed by 45 min incubation with 55 mM iodacetamide at room temperature. In-gel tryptic digestion (Sequencing Grade Modified trypsin, Promega) was performed overnight at 37 °C and peptide extraction was carried out by consecutive incubations with 0.1% trifluoroacetic acid in 60% acetonitrile. Extracted peptides were vacuum-dried and resuspended in 0.1% formic acid.

Nano-HPLC was coupled to a linear ion trap mass spectrometer (LTQ Velos, Thermo Scientific). Nano ESI-source parameters were set as follows: spray voltage 2.3 kV and capillary temperature 260 °C. The equipment was operated in a data dependent acquisition mode: a full MS scan acquired in positive ion mode (m/z between 300 and 1800 Da) was followed by fragmentation of the 10 most intense ions (normalized collision energy: 35, activation Q: 0.25; activation time: 15 ms) using a dynamic exclusion list.

### Bioinformatics analyses

Bioinformatics data analysis was performed using PatternLab for Proteomics V5 (http://patternlabforproteomics.org) (48). A target-decoy database was generated using *M. smegmatis* strain ATCC 700084/MC2155 sequences (downloaded from Uniprot on 2016-03-29) plus the sequence of the Strep-tagged FhaA and 127 most common mass spectrometry contaminants. Search parameters were set as follows: tryptic peptides; oxidation of Met as variable modification; carbamidomethylation as fixed Cys modification; 800 ppm of tolerance from the measured precursor m/z. Search Engine Processor (SEPro) was used to filter peptide spectrum matches to an FDR< 1% at the protein level. Identification of proteins exclusively detected in FhaA purified crosslinked complexes was performed using PatternLab’s Approximately Area Proportional Venn Diagram module. Proteins present in at least four out of five *Msmeg_fhaA* replicates, but absent in all control replicates, and statistically validated using the Bayesian model integrated into PatternLab for proteomics Venn Diagram module were considered part of FhaA interactome. In addition, TFold module was used to pinpoint proteins statistically enriched in FhaA complexes according to their spectral counts (BH q value: 0.05, F Stringency: 0.04 and L-stringency: 0.6) (49,50).

### Cell surface hydrophobicity test

Surface hydrophobicity was quantified using the Microbial Adhesion to Hydrocarbon (MATH) method (51). For that purpose, cells were partitioned using a two-phase system, according to previous reports (52). Exponential growth phase cultures were washed and resuspended in PBS until a final OD600 of 0.7. Samples were mixed with xylene in a 1:1 ratio and incubated 15 min at room temperature to allow partitioning. OD600 of aqueous layer was determined. Hydrophobicity index represents the percentage of initial aqueous layer absorbance retained in the xylene fraction after partitioning. Experiments were performed by triplicate.

### Biofilm formation assay

Microtiter dish biofilm formation assay was performed as previously described (47). Briefly, bacterial cultures were loaded in 96 wells plates to an initial OD600 of 0.1. Biofilm formation was evaluated by biological triplicates when static cultures of both strains reached identical OD600. Staining was performed with crystal violet. Biofilms were destained in 30% acetic acid and OD570 of the retained crystal violet was determined. Experiments were performed in triplicate.

### Growth curve

To ensure that observed differences in biofilm formation are not due to differences in biomass, a static growth curve was recorded, with cells grown in the same conditions used for biofilm assays. Briefly, 96 wells plates were loaded with three independent cultures of each strain in quintuplicate and incubated at room temperature without shaking. OD600 was measured once a day for 10 days.

### Fluorescence microscopy and Image acquisition and analysis

Exponential growth phase cultures were incubated with 50 μM HADA for 30 min. Samples were loaded into glass slides and allowed to dry at 37 °C. Sulforhodamine-DHPE 10 μg/mL was added and allowed to dry. Slides were washed with sterile water and mounted in 10% BSA. Images were acquired with a Zeiss LSM 880 confocal laser scanning microscope, equipped with a plan-apochromatic 63x/1.4 oil immersion objective. Image acquisition was performed in channel mode with a pixel size of 0.105 μm and a resolution of 256×256. HADA excitation was performed with a 405 nm laser and emitted light was collected in the range between 415 and 480 nm; Sulforhodamine-DHPE excitation was performed using a 561 nM laser and light emission was collected between 580 and 620 nm. mScarlet excitation was performed using a 561 nm laser and emitted light was collected in the range between 570 and 655 nm.

Images were processed and analysed using Image J (53). All cell length measurements were performed using the Sulforhodamine-DHPE signal; all calculated parameters were obtained from the HADA or mScarlet-FhaA signals.

Control HADA foci were detected using *Find maxima tool* of Image J. Prominence was set to detect 2 (non-septate) or 3 (septate) foci per cell in control strain. The same setting was then applied to *Msmeg*_*fhaA* strain. For comparative purposes, and to account for differences in cell length, distances between HADA foci were expressed as a fraction of the total length.

For WT, *Msmeg*_*ΔfhaA* and *Msmeg_ΔfhaA*_*fhaA* analysis, we use ImageJ to obtain the intensity profile of HADA drawing a segmented line across the longitudinal axis. The line has a width of 10 pixels, corresponding to approximately 1.06 μm, and a spatial resolution of 0.09μm. Each profile is computed as the average intensity across the line width, and cell length was normalized. For each strain, we compute a representative profile by averaging all the bacteria profiles in the strain. Also, we obtained the position and intensity of septum and poles for each bacterium in the strain by using *findpeaks* Scipy function. Septum position was ultimately utilized to select and align both the old and new poles, with the new pole positioned closest to the septum. To determine the pole intensity decay towards the septum we consider 10 points of the profile curve and fit it to a 1st order polynomial. We consider the slope of the fit curve as the value representing the decay.

For the *Msmeg_mscarlet_fhaA* analysis, ImageJ was used to extract the HADA fluorescence profile, following the procedure described above. Profiles of each individual bacterium were length-normalized and aligned according to their type of pole (old or new). This classification allowed us to determine the fluorescence intensity of mScarlet-FhaA associated with each type of pole.

Statistical comparisons between strains were performed using one-way ANOVA or T-test for normally distributed data, and Kolmogorov-Smirnov (2 samples) or Kruskal-Wallis (3 samples) for not normally distributed data.

### Sample preparation for transmission electron microscopy

For TEM analysis, samples were fixed with 2.5% glutaraldehyde and 4% formaldehyde in 0.1 M cacodylate buffer (pH 7.2) for 2 h and post-fixed for 1 h in 1% OsO_4_ with 2.5% potassium ferrocyanide in the same buffer. Samples were then dehydrated in acetone and embedded in Polybed 812 resin (Polysciences). Ultrathin (60 nm) sections were stained with 5% uranyl acetate (40 min) and 2% lead citrate (5 min) before observation using a JEOL 1200 EX transmission electron microscope at 120 kV equipped with a camera Megaview G2 CCD 1k.

### Electron tomography

As previously established for electron tomography (54–56) samples processed for TEM were sectioned (200 nm thick serial sections) in a PowerTome XL ultramicrotome (RMC Boeckeler) and collected onto formvar-coated copper slot grids, then stained with 5% (w/v) uranyl acetate and lead citrate. In addition, 10 nm colloidal gold particles (Gold colloid, Sigma-Aldrich) were used as fiducial markers during the tilted series’ alignment. Finally, a single-axis tilt series (± 65° with 2° increment) was collected from the samples using a Tecnai G2 F20 transmission electron microscope (Thermo Fisher Scientific) operating at 200 kV in TEM mode with a camera AMT CMOS 4K. Tomographic tilt series were processed using IMOD version 4.9.13 (University of Colorado, USA). Projections were aligned by cross-correlation. The final alignment was performed using 10 nm fiducial gold particles followed by weighted back-projection reconstruction. Manual segmentation, surface rendering and the thickness map analysis were performed with the Amira software (Thermo Fisher Scientific).

### Scanning electron microscopy

Samples were fixed with 2.5% glutaraldehyde and 4% formaldehyde in 0.1 M cacodylate buffer (pH 7.2) for 2 h and then adhered to poly-L-lysine treated coverslips. Next, the coverslips were washed with 0.1 M sodium cacodylate buffer and post-fixed for 40 min in 1% OsO4 with 2.5% potassium ferrocyanide. After another washing cycle of three rounds, the samples were dehydrated through a series of increasing concentrations (30–100%) of ethanol. Finally, the samples were critical-point-dried in liquid CO2 in a Leica EM CPD300 apparatus and sputtering with a 2-nm-thick platinum coat in a Quorum Q150V Plus apparatus. Samples were observed using a FEG Quattro S scanning electron microscope (Thermo Fisher Scientific) operating at 5 kV.

### Morphometry

Cell wall thickness measurements were carried out on images obtained from ultra-thin TEM sections, whereas cell width measurements were derived from SEM images using the Fiji/Image J software. Two opposing regions of each cell were assessed for thickness, while three regions (ends and center) of each cell were measured for width. The mean values were calculated based on data obtained from 30 cells in each experimental group. Statistical analyses were conducted using the Kolmogorov Smirnov (TEM) or one-way ANOVA (SEM), with significance set at p < 0.05.

### LAURDAN staining, image acquisition and spectral phasor analysis

Exponential growth phase cultures (OD 600 0.8) were centrifuged and washed in PBS. Pellets were resuspended in 50 μl of 0.05 mM LAURDAN-DMSO in PBS and incubated at 37° C and 220 rpm for 2 h. Live bacteria were mounted in agarose patches and visualized using a Zeiss LSM 880 spectral confocal laser scanning microscope, equipped with a plan-apochromatic 63x/1.4 oil immersion objective. LAURDAN excitation was performed in lambda mode, using a 405 nm laser for excitation and emission was collected in the range from 418-718 nm, in 30 channels, 10 nm each, and an extra channel for transmitted light. Images were acquired with a 256 x 256-pixel resolution and a scan zoom of 10x (pixel size 0.05 x 0.05 μm; pixel time 0.67μs). As LAURDAN emission spectrum is sensitive to the lipid composition and dipolar relaxation, it may be used to assess water accessibility in the environment in which the probe is embedded. Spectral phasor analysis of LAURDAN emission was performed using FLIM module of SimFCS 4 software (www.lfd.uci.edu/globals). Briefly, LAURDAN emission spectra from each pixel were Fourier transformed and, G and S, (corresponding to the real and imaginary parts of the first harmonic of Fourier transform) were used as x and y coordinates of the phasor plot. Pixels with similar spectral properties cluster together on the plot. While angular position (Φ) of clustered pixels into the phasor plot provides information about the emission-spectra-center-of-mass, spectral widening relies on radial position (M). Each pixel of the image is associated with a phasor in the phasor plot, and each phasor maps to pixels in the image.

## Data Availability

The mass spectrometry interactomics data have been deposited to the ProteomeXchange Consortium via the PRIDE partner repository with the dataset identifier PXD054354.

## Supplementary Figures

**Figure S1. Static growth curve.**

Cells were grown in 7H9 medium at 37 °C without agitation. Assay was performed in 96 wells plates, using three independent cultures of each strain in quintuplicates. DO 600 was measured each day for 10 days. Note that both strains reach the same optical density at stationary phase.

**Figure S2**

**A)** Transmission electron microscopy (TEM) images of ultrathin sections from control and *Msmeg*_*fhaA* strains, showing differences on cell wall thickness and appearance. The asterisks (*) highlight areas of abnormal cell wall increase observed for *Msmeg*_*fhaA*. This strain has nearly twice the cell wall thickness compared to control strain (Figure 2C). Scale bar: 200 nm. B) Histograms representing the cell wall thickness distribution measured in tomograms (Figure 2D) for control and *Msmeg*_*fhaA*. n=30.

**Figure S3.**

**A)** Alignment of average HADA fluorescence profiles of Figure 6 B between strains. Average profiles are length normalized for comparative purposes. Poles 1 and 2, corresponding to fast and slow growing pole, respectively, are indicated. **B)** Box plots showing comparison of slopes of fluorescence intensity corresponding to HADA incorporation from the tip between strains for each pole, labelled as pole 1 and 2. Both poles of *Msmeg*_Δ*fhaA* have statistically significant smaller slopes than pole 1 and pole 2 from WT and *Msmeg*_Δ*fhaA*_*fhaA.* * Indicates statistically significant difference determined by Kolmogorov-Smirnov test, p < 0.05; n >100 cells for each group. Box represents 25%-75% percentile and median; values outside whiskers represents outliers.

## Acknowledgments

We express our gratitude to Magdalena Portela for the excellent technical support and to Dr. Adriana Parodi for her insightful discussions on the results. We also thank Dr. Raghunand Tirumalai for kindly providing us with the mc^2^6 strains (*Msmeg_ΔfhaA; Msmeg_ΔfhaA_fhaA* and WT). We extend our gratitude to Dr. Martin Graña for his critical review and valuable contributions to improving the manuscript and BSc. Bruno Schuty for fruitful discussions.

This work was funded by grants from the Agencia Nacional de Investigación e Innovación, Uruguay (FCE_1_2014_1_104045) (RD), FOCEM (MERCOSUR Structural Convergence Fund, COF 03/11) and ECOS-Sud France-Uruguay (contract U20B02, RD and AMW), the Agence Nationale de la Recherche (ANR,France), contract ANR-18-CE11-0017 (P.M.A.), and by institutional grants from the Institut Pasteur, the CNRS, and Université Paris Cité. MG, JR, BR and ART were supported by a fellowship from ANII [POS_NAC_2012_1_8824, POS_NAC_2015_1_109755, POS_FCE_2015_1_1005186 and POS_FCE_2020 1009183]. JR and ART were supported by the Comisión Académica de Posgrado, UdelaR, Uruguay. MG, JR, BR and ART were supported by PEDECIBA.

## References

1. Bagcchi S. WHO’s Global Tuberculosis Report 2022. Lancet Microbe. 2023 Jan;4(1):e20.

2. Kieser KJ, Rubin EJ. How sisters grow apart: mycobacterial growth and division. Nat Rev Microbiol. 2014 Aug;12(8):550–62.

3. Brennan PJ. Structure, function, and biogenesis of the cell wall of Mycobacterium tuberculosis. Tuberc Edinb. 2003;83(1–3):91–7.

4. Jarlier V, Nikaido H. Mycobacterial cell wall: Structure and role in natural resistance to antibiotics. FEMS Microbiol Lett. 1994 Oct;123(1–2):11–8.

5. Meniche X, Otten R, Siegrist MS, Baer CE, Murphy KC, Bertozzi CR, et al. Subpolar addition of new cell wall is directed by DivIVA in mycobacteria. Proc Natl Acad Sci. 2014;111(31):E3243–51.

6. Aldridge BB, Fernandez-Suarez M, Heller D, Ambravaneswaran V, Irimia D, Toner M, et al. Asymmetry and Aging of Mycobacterial Cells Lead to Variable Growth and Antibiotic Susceptibility. Science. 2012 Jan 6;335(6064):100–4.

7. Donovan C, Bramkamp M. Cell division in Corynebacterineae. Front Microbiol. 2014;5:132.

8. Bellinzoni M, Wehenkel AM, Duran R, Alzari PM. Novel mechanistic insights into physiological signaling pathways mediated by mycobacterial Ser/Thr protein kinases. Microbes Infect. 2019 Jul;21(5–6):222–9.

9. Sureka K, Hossain T, Mukherjee P, Chatterjee P, Datta P, Kundu M, et al. Novel role of phosphorylation-dependent interaction between FtsZ and FipA in mycobacterial cell division. PLoS One. 2010 Jan 6;5(1):e8590.

10. Gee CL, Papavinasasundaram KG, Blair SR, Baer CE, Falick AM, King DS, et al. A phosphorylated pseudokinase complex controls cell wall synthesis in mycobacteria. Sci Signal. 2012 Jan 24;5(208):ra7.

11. Fernandez P, Saint-Joanis B, Barilone N, Jackson M, Gicquel B, Cole ST, et al. The Ser/Thr protein kinase PknB is essential for sustaining mycobacterial growth. J Bacteriol. 2006 Nov;188(22):7778–84.

12. Wehenkel A, Bellinzoni M, Grana M, Duran R, Villarino A, Fernandez P, et al. Mycobacterial Ser/Thr protein kinases and phosphatases: physiological roles and therapeutic potential. Biochim Biophys Acta. 2008 Jan;1784(1):193–202.

13. Roumestand C, Leiba J, Galophe N, Margeat E, Padilla A, Bessin Y, et al. Structural insight into the Mycobacterium tuberculosis Rv0020c protein and its interaction with the PknB kinase. Structure. 2011 Oct 12;19(10):1525–34.

14. Turapov O, Forti F, Kadhim B, Ghisotti D, Sassine J, Straatman-Iwanowska A, et al. Two Faces of CwlM, an Essential PknB Substrate, in Mycobacterium tuberculosis. Cell Rep. 2018 Oct 2;25(1):57–67 e5.

15. Viswanathan G, Yadav S, Joshi SV, Raghunand TR. Insights into the function of FhaA, a cell division-associated protein in mycobacteria. FEMS Microbiol Lett. 2017 Jan 1;364(2).

16. Lougheed KE, Bennett MH, Williams HD. An in vivo crosslinking system for identifying mycobacterial protein-protein interactions. J Microbiol Methods. 2014 Oct;105:67–71.

17. Gupta KR, Gwin CM, Rahlwes KC, Biegas KJ, Wang C, Park JH, et al. An essential periplasmic protein coordinates lipid trafficking and is required for asymmetric polar growth in mycobacteria. Elife. 2022 Nov 8;11.

18. Fay A, Czudnochowski N, Rock JM, Johnson JR, Krogan NJ, Rosenberg O, et al. Two Accessory Proteins Govern MmpL3 Mycolic Acid Transport in Mycobacteria. mBio. 2019 Jun 25;10(3).

19. Plocinski P, Arora N, Sarva K, Blaszczyk E, Qin H, Das N, et al. Mycobacterium tuberculosis CwsA interacts with CrgA and Wag31, and the CrgA-CwsA complex is involved in peptidoglycan synthesis and cell shape determination. J Bacteriol. 2012 Dec;194(23):6398–409.

20. Mir M, Prisic S, Kang CM, Lun S, Guo H, Murry JP, et al. Mycobacterial gene cuvA is required for optimal nutrient utilization and virulence. Infect Immun. 2014;82(10).

21. Belardinelli JM, Stevens CM, Li W, Tan YZ, Jones V, Mancia F, et al. The MmpL3 interactome reveals a complex crosstalk between cell envelope biosynthesis and cell elongation and division in mycobacteria. Sci Rep. 2019 Jul 24;9(1):10728.

22. Plocinski P, Martinez L, Sarva K, Plocinska R, Madiraju M, Rajagopalan M. Mycobacterium tuberculosis CwsA overproduction modulates cell division and cell wall synthesis. Tuberculosis. 2013 Dec;93:S21–7.

23. Gurcha SS, Baulard AR, Kremer L, Locht C, Moody DB, Muhlecker W, et al. Ppm1, a novel polyprenol monophosphomannose synthase from Mycobacterium tuberculosis. Biochem J. 2002 Jul 15;365(Pt 2):441–50.

24. Rana AK, Singh A, Gurcha SS, Cox LR, Bhatt A, Besra GS. Ppm1-Encoded Polyprenyl Monophosphomannose Synthase Activity Is Essential for Lipoglycan Synthesis and Survival in Mycobacteria. Tyagi AK, editor. PLoS ONE. 2012 Oct 31;7(10):e48211.

25. Cashmore TJ, Klatt S, Yamaryo-Botte Y, Brammananth R, Rainczuk AK, McConville MJ, et al. Identification of a Membrane Protein Required for Lipomannan Maturation and Lipoarabinomannan Synthesis in Corynebacterineae. J Biol Chem. 2017 Mar 24;292(12):4976–86.

26. Lee JH, Jeong H, Kim Y, Lee HS. Corynebacterium glutamicum whiA plays roles in cell division, cell envelope formation, and general cell physiology. Antonie Van Leeuwenhoek. 2020;113:629–41.

27. Pickford H, Alcock E, Singh A, Kelemen G, Bhatt A. A mycobacterial DivIVA domain-containing protein involved in cell length and septation. Microbiology. 2020 Sep 1;166(9):817–25.

28. Chakraborty P, Kumar A. The extracellular matrix of mycobacterial biofilms: could we shorten the treatment of mycobacterial infections? Microb Cell. 2019 Jan 18;6(2):105–22.

29. Mazumder S, Falkinham JO, Dietrich AM, Puri IK. Role of hydrophobicity in bacterial adherence to carbon nanostructures and biofilm formation. Biofouling. 2010 Jan 13;26(3):333–9.

30. Malacrida L, Jameson DM, Gratton E. A multidimensional phasor approach reveals LAURDAN photophysics in NIH-3T3 cell membranes. Sci Rep. 2017 Aug 23;7(1):9215.

31. Malacrida L, Gratton E. LAURDAN fluorescence and phasor plots reveal the effects of a H2O2 bolus in NIH-3T3 fibroblast membranes dynamics and hydration. Free Radic Biol Med. 2018 Nov 20;128:144–56.

32. Kuru E, Tekkam S, Hall E, Brun YV, Van Nieuwenhze MS. Synthesis of fluorescent D-amino acids and their use for probing peptidoglycan synthesis and bacterial growth in situ. Nat Protoc. 2015;10(1):33–52.

33. Nguyen L, Scherr N, Gatfield J, Walburger A, Pieters J, Thompson CJ. Antigen 84, an Effector of Pleiomorphism in *Mycobacterium smegmatis*. J Bacteriol. 2007 Nov;189(21):7896–910.

34. Joyce G, Williams KJ, Robb M, Noens E, Tizzano B, Shahrezaei V, et al. Cell Division Site Placement and Asymmetric Growth in Mycobacteria. Driks A, editor. PLoS ONE. 2012 Sep 10;7(9):e44582.

35. Hannebelle MTM, Ven JXY, Toniolo C, Eskandarian HA, Vuaridel-Thurre G, McKinney JD, et al. A biphasic growth model for cell pole elongation in mycobacteria. Nat Commun. 2020 Jan 23;11(1):452.

36. Wu KJ, Zhang J, Baranowski C, Leung V, Rego EH, Morita YS, et al. Characterization of Conserved and Novel Septal Factors in Mycobacterium smegmatis. J Bacteriol. 2018 Mar 15;200(6).

37. Kavunja HW, Biegas KJ, Banahene N, Stewart JA, Piligian BF, Groenevelt JM, et al. Photoactivatable Glycolipid Probes for Identifying Mycolate–Protein Interactions in Live Mycobacteria. J Am Chem Soc. 2020 Apr 29;142(17):7725–31.

38. Bou Raad R, Méniche X, De Sousa-d’Auria C, Chami M, Salmeron C, Tropis M, et al. A Deficiency in Arabinogalactan Biosynthesis Affects *Corynebacterium glutamicum* Mycolate Outer Membrane Stability. J Bacteriol. 2010 Jun;192(11):2691–700.

39. Kang CM, Nyayapathy S, Lee JY, Suh JW, Husson RN. Wag31, a homologue of the cell division protein DivIVA, regulates growth, morphology and polar cell wall synthesis in mycobacteria. Microbiology. 2008 Mar;154(Pt 3):725–35.

40. Kieser KJ, Boutte CC, Kester JC, Baer CE, Barczak AK, Meniche X, et al. Phosphorylation of the Peptidoglycan Synthase PonA1 Governs the Rate of Polar Elongation in Mycobacteria. Behr MA, editor. PLOS Pathog. 2015 Jun 26;11(6):e1005010.

41. Jani C, Eoh H, Lee JJ, Hamasha K, Sahana MB, Han JS, et al. Regulation of Polar Peptidoglycan Biosynthesis by Wag31 Phosphorylation in Mycobacteria. BMC Microbiol. 2010 Dec;10(1):327.

42. Chung ES, Johnson WC, Aldridge BB. Types and functions of heterogeneity in mycobacteria. Nat Rev Microbiol. 2022 Sep;20(9):529–41.

43. Gil M, Lima A, Rivera B, Rossello J, Urdaniz E, Cascioferro A, et al. New substrates and interactors of the mycobacterial Serine/Threonine protein kinase PknG identified by a tailored interactomic approach. J Proteomics. 2019 Feb 10;192:321–33.

44. Bindels DS, Haarbosch L, Van Weeren L, Postma M, Wiese KE, Mastop M, et al. mScarlet: a bright monomeric red fluorescent protein for cellular imaging. Nat Methods. 2017;14(1):53–6.

45. Martinez M, Petit J, Leyva A, Sogues A, Megrian D, Rodriguez A, et al. Eukaryotic-like gephyrin and cognate membrane receptor coordinate corynebacterial cell division and polar elongation. Nat Microbiol. 2023 Sep 7;8(10):1896–910.

46. Sogues A, Martinez M, Gaday Q, Ben Assaya M, Graña M, Voegele A, et al. Essential dynamic interdependence of FtsZ and SepF for Z-ring and septum formation in Corynebacterium glutamicum. Nat Commun. 2020;11(1).

47. Rossello J, Lima A, Gil M, Rodriguez Duarte J, Correa A, Carvalho PC, et al. The EAL-domain protein FcsR regulates flagella, chemotaxis and type III secretion system in Pseudomonas aeruginosa by a phosphodiesterase independent mechanism. Sci Rep. 2017 Aug 31;7(1):10281.

48. Santos MDM, Lima DB, Fischer JSG, Clasen MA, Kurt LU, Camillo-Andrade AC, et al. Simple, efficient and thorough shotgun proteomic analysis with PatternLab V. Nat Protoc. 2022;17(7).

49. Carvalho PC, Lima DB, Leprevost FV, Santos MD, Fischer JS, Aquino PF, et al. Integrated analysis of shotgun proteomic data with PatternLab for proteomics 4.0. Nat Protoc. 2016 Jan;11(1):102–17.

50. Carvalho PC, Yates JR, Barbosa VC. Improving the TFold test for differential shotgun proteomics. Bioinformatics. 2012 Jun 15;28(12):1652–4.

51. Rosenberg M, Gutnick D, Rosenberg E. Adherence of bacteria to hydrocarbons: A simple method for measuring cell-surface hydrophobicity. FEMS Microbiol Lett. 1980 Sep;9(1):29–33.

52. Rosenberg M. Microbial adhesion to hydrocarbons: twenty-five years of doing MATH. FEMS Microbiol Lett. 2006 Sep;262(2):129–34.

53. Schindelin J, Rueden CT, Hiner MC, Eliceiri KW. The ImageJ ecosystem: An open platform for biomedical image analysis. Mol Reprod Dev. 2015 Jul;82(7–8):518– 29.

54. Girard-Dias W, Alcântara CL, Cunha-e-Silva N, de Souza W, Miranda K. On the ultrastructural organization of Trypanosoma cruzi using cryopreparation methods and electron tomography. Histochem Cell Biol. 2012;138:821–31.

55. Wendt C, Rachid R, de Souza W, Miranda K. Electron tomography characterization of hemoglobin uptake in Plasmodium chabaudi reveals a stage-dependent mechanism for food vacuole morphogenesis. J Struct Biol. 2016;194(2):171–9.

56. Girard-Dias W, Augusto I, VA Fernandes T, G. Pascutti P, de Souza W, Miranda K. A spatially resolved elemental nanodomain organization within acidocalcisomes in Trypanosoma cruzi. Proc Natl Acad Sci. 2023;120(16):e2300942120.

57. Bateman A, Martin MJ, Orchard S, Magrane M, Ahmad S, Alpi E, et al. UniProt: the Universal Protein Knowledgebase in 2023. Nucleic Acids Res. 2023;51(D1).

58. Hallgren J, Tsirigos KD, Pedersen MD, Juan J, Armenteros A, Marcatili P, et al. DeepTMHMM predicts alpha and beta transmembrane proteins using deep neural networks. bioRxiv. 2022;

59. Hayashi JM, Luo CY, Mayfield JA, Hsu T, Fukuda T, Walfield AL, et al. Spatially distinct and metabolically active membrane domain in mycobacteria. Proc Natl Acad Sci. 2016 May 10;113(19):5400–5.

60. Zhu J, Wolf ID, Dulberger CL, Won HI, Kester JC, Judd JA, et al. Spatiotemporal localization of proteins in mycobacteria. Cell Rep. 2021 Dec;37(13):110154.

