## Supplementary figures and images for "FhaA plays a key role in mycobacterial polar elongation and asymmetric growth"

### Supplementary Figures 1-3

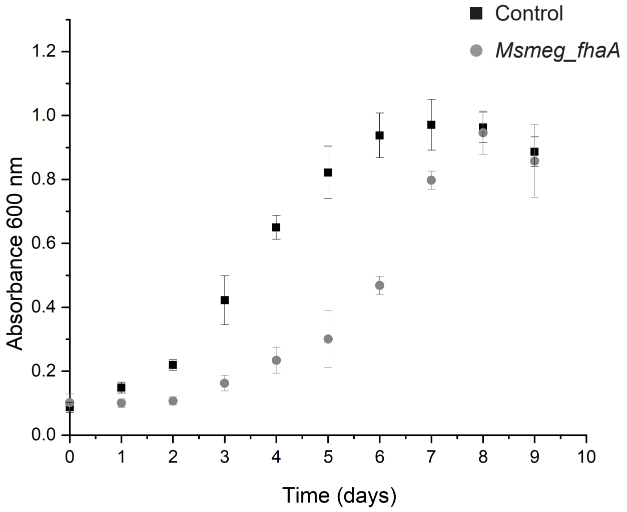

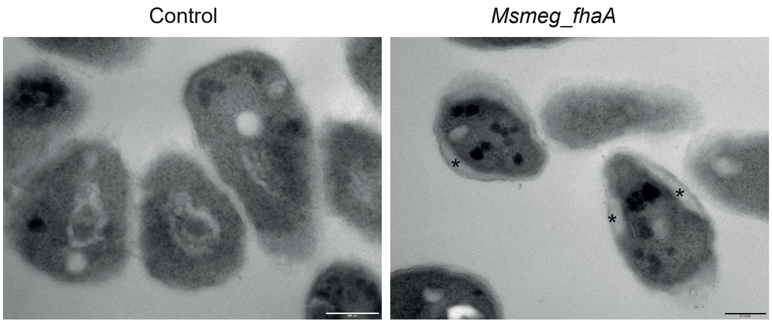

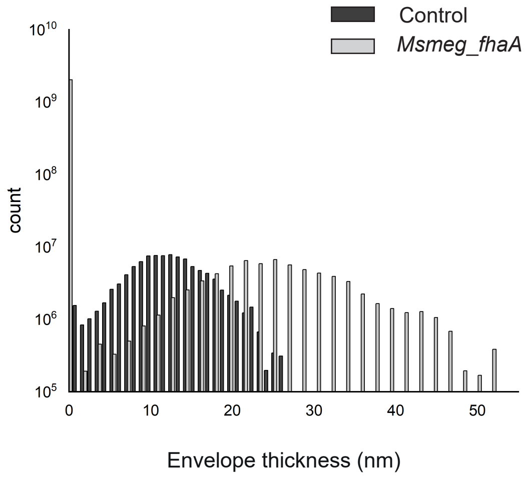

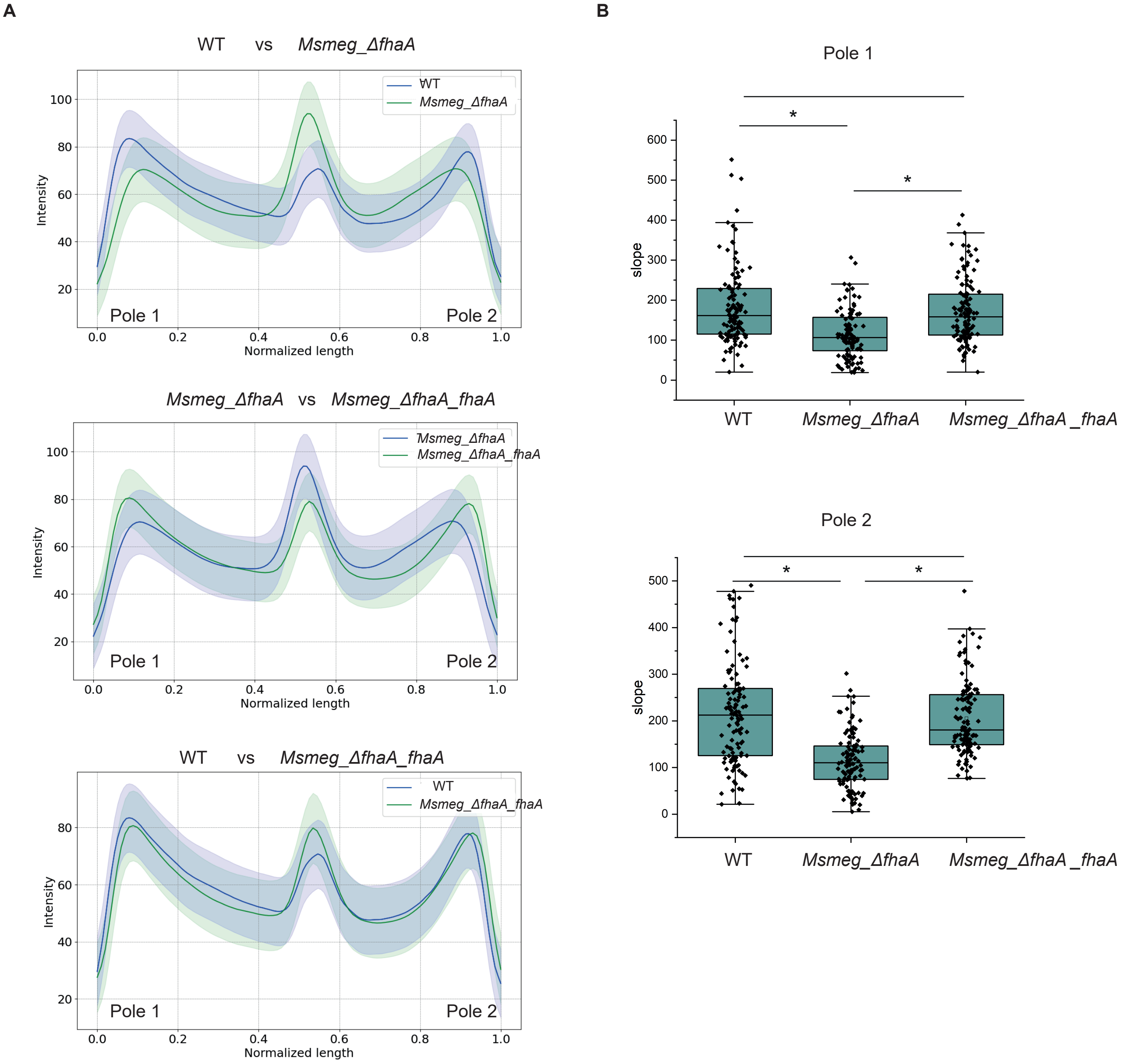
